# Host glycans trap bacterial lectins to trigger NLRP3 inflammasome activation

**DOI:** 10.64898/2026.11.24.768555

**Authors:** Shengnan Zou, Olga Makshakova, Xin Wen, Svenja Wöhrle, Simon Lagies, Shuxin Shi, Clara Dufossez, Nele Horn, Lena Wolter, Bernd Kammerer, Olaf Groß, Daniel Fisch, Winfried Römer

## Abstract

*Pseudomonas aeruginosa* poses a severe threat to immunocompromised patients through its multifaceted virulence arsenal. Although diverse bacterial components activate the NLRP3 inflammasome, the precise molecular triggers driving this process remain largely elusive. Here, we reveal that host surface glycans effectively trap the *P. aeruginosa* lectins LecA and LecB, initiating carbohydrate-dependent inflammasome activation. This lectin-mediated cascade exhibits striking host-species and cell-type specificity. The L-fucose-specific LecB activates human blood monocytes and THP-1 cells, where glycosylated β1-integrin contributes to signal transduction, whereas the D-galactose-specific LecA is more potent in murine bone marrow-derived macrophages and dendritic cells. This mechanism requires potassium efflux and displays cell-specific sensitivity to reactive oxygen species. Infection with live bacteria recapitulates the lectin-mediated inflammasome response. Collectively, our findings unveil an unexpected role for host glycans as critical, previously overlooked checkpoints gating *P. aeruginosa*-driven inflammation, offering fresh insights into host-pathogen dynamics and exposing novel therapeutic targets.

## INTRODUCTION

*Pseudomonas aeruginosa* (*P. aeruginosa*) is a highly adaptable Gram-negative opportunistic bacterium that causes severe acute and chronic infections, particularly in immunocompromised individuals and hospitalized patients.^1,2^ Its intrinsic antibiotic resistance and ability to persist in hostile environments, including hypoxic and nutrient-limited niches, make it a major clinical challenge.^3–5^ Reflecting this threat, the World Health Organization recently classified carbapenem-resistant *P. aeruginosa* as a high-priority pathogen, underscoring the urgent need to better understand mechanisms of pathogenesis and host damage.^6^

This versatility is rooted in a broad arsenal of virulence factors that enable adhesion, invasion, immune modulation and long-term persistence.^5^ Surface components such as lipopolysaccharide (LPS), type IV pili, and flagella facilitate host interaction and motility,^4,7–10^ while secretion systems, most notably the type III secretion system (T3SS), inject effector proteins that reprogram host signaling.^10–12^ In parallel, secreted exopolysaccharides promote biofilm formation and antibiotic tolerance,^13^ quorum-sensing networks coordinate virulence in a cell density-dependent manner.^14,15^

Among these virulence factors, the carbohydrate-binding lectins LecA (PA-IL) and LecB (PA-IIL) have emerged as key mediators of host-pathogen interactions.^16^ Both form homo-tetramers, with each monomer (12.8 kDa for LecA; 11.7 kDa for LecB) containing one carbohydrate-binding site.^17,18^ LecA has a high affinity for D-galactose and binds the host glycosphingolipid globotriaosylceramide (Gb3, also known as CD77). This interaction induces Gb3 clustering and signaling processes that drive membrane engulfment and bacterial uptake via the so-called lipid-zipper mechanism.^19–24^ Conversely, host cells try to restrict bacterial penetration by recruiting septins, which assemble into rigid barriers.^25^

LecB preferentially binds to L-fucose and, to a lesser extent, D-mannose.^26–30^ Beyond stabilizing *P. aeruginosa* biofilms via binding to the exopolysaccharide Psl,^31^ LecB disrupts epithelial integrity through multiple mechanisms. In human lung epithelial cells, it promotes β-catenin degradation and impairs tissue repair,^32^ while in keratinocytes, it binds IGF-1R and induces receptor internalization without activation, causing endocytic misrouting and suppressed growth factor signaling.^33^ In MDCK cells, LecB triggers the internalization of integrins, thereby impairing epithelial wound healing.^34^ In the same cells, it activates Src-PI3K/Akt signaling and recruits caveolin-1 to promote *P. aeruginosa* invasion from the apical side.^35^ Beyond these effects, LecB modulates host immunity by impairing dendritic cell migration and T cell activation in vivo, an effect that is reversed by LecB glycomimetic inhibitors.^36^ However, the extent to which *P. aeruginosa* lectins target other critical components of host innate immunity, particularly macrophages, remains to be fully elucidated.

The innate immune system constitutes the first line of defense against bacterial pathogens using germline-encoded pattern recognition receptors (PRRs) to detect pathogen- and damage-associated molecular patterns (PAMPs, DAMPs).^37–40^ In the cytosol, specific PRRs such as NOD-like receptors (NLR) and AIM2, assemble into multiprotein complexes called inflammasomes, which are broadly classified into canonical and non-canonical pathways. Canonical inflammasomes comprise a receptor (e.g., NLRP1, NLRP3, NLRC4, AIM2), the adaptor ASC (apoptosis-associated speck-like protein containing a CARD) in many cases, and the effector caspase-1. Their activation triggers proteolytic activation of caspase-1, which promotes the maturation of IL-1β and cleaves gasdermin D (GSDMD) to induce pyroptosis, a pro-inflammatory form of cell death. In contrast, non-canonical inflammasomes are mediated by murine caspase-11 and human caspase-4/5, which directly sense cytosolic LPS and trigger pyroptosis through GSDMD cleavage.^41–45^ Among these, the NLRP3 inflammasome is the most broadly responsive and extensively studied.^46,47^

While *P. aeruginosa* employs several bacterial components to trigger the NLRP3 inflammasome, such as outer membrane vesicles and T3SS-associated signals,^48,49^ the precise molecular drivers behind this response remain an active area of discovery. Here, we demonstrate that host glycans trap the *P. aeruginosa* lectins LecA and LecB, triggering carbohydrate-dependent activation of the NLRP3 inflammasome. Furthermore, we show that this lectin-mediated response is specific to both host species and cell types. Our findings unveil an unexpected role for host glycans, establishing them as a critical, hitherto unrecognized checkpoint in lectin-driven inflammasome activation, and fundamentally reshaping our understanding of the host-pathogen interface.

## RESULTS

### LecB triggers carbohydrate-dependent inflammasome activation in human macrophages

To study lectin-driven inflammasome activation, we used PMA-differentiated THP-1 cells as a well-established in vitro model for human macrophages.^50–52^ Immunoblot analysis of LPS-primed and LecB-stimulated THP-1 cells revealed a time-dependent response: After 24 hours, LecB induced robust release of cleaved caspase-1 (p20) and mature IL-1β (p17) into the supernatant, along with GSDMD-N (p35) in cell lysates, comparable to the canonical NLRP3 inflammasome agonist Nigericin as control. These cleavage products were absent at 3 hours and only weakly detectable at 6 hours (Figure 1A). ELISA confirmed a progressive increase in IL-1β secretion, with slight elevation at 3 hours and further increases at 6 hours, showing a LecB concentration-dependent effect at both time points, whereas LPS-alone had no effect (Figure 1B). By 24 hours, IL-1β secretion was maximal, and levels at 0.5 and 1 μM LecB had converged. Therefore, we chose to use 0.5 μM LecB for 24 hours in subsequent experiments. In comparison and as expected, Nigericin induced rapid, near-maximal IL-1β release at 3 hours, with no further increase at later time points, whereas LecB reached comparable levels by 24 hours (Figure 1B). In contrast, the galactose-binding lectin LecA did not induce caspase-1 activation, GSDMD cleavage, or IL-1β release in THP-1 cells (Figure S1A). Consistently, ELISA showed no significant increase in IL-1β secretion compared to the LPS-primed controls (Figure S1B).

**Figure 1.**
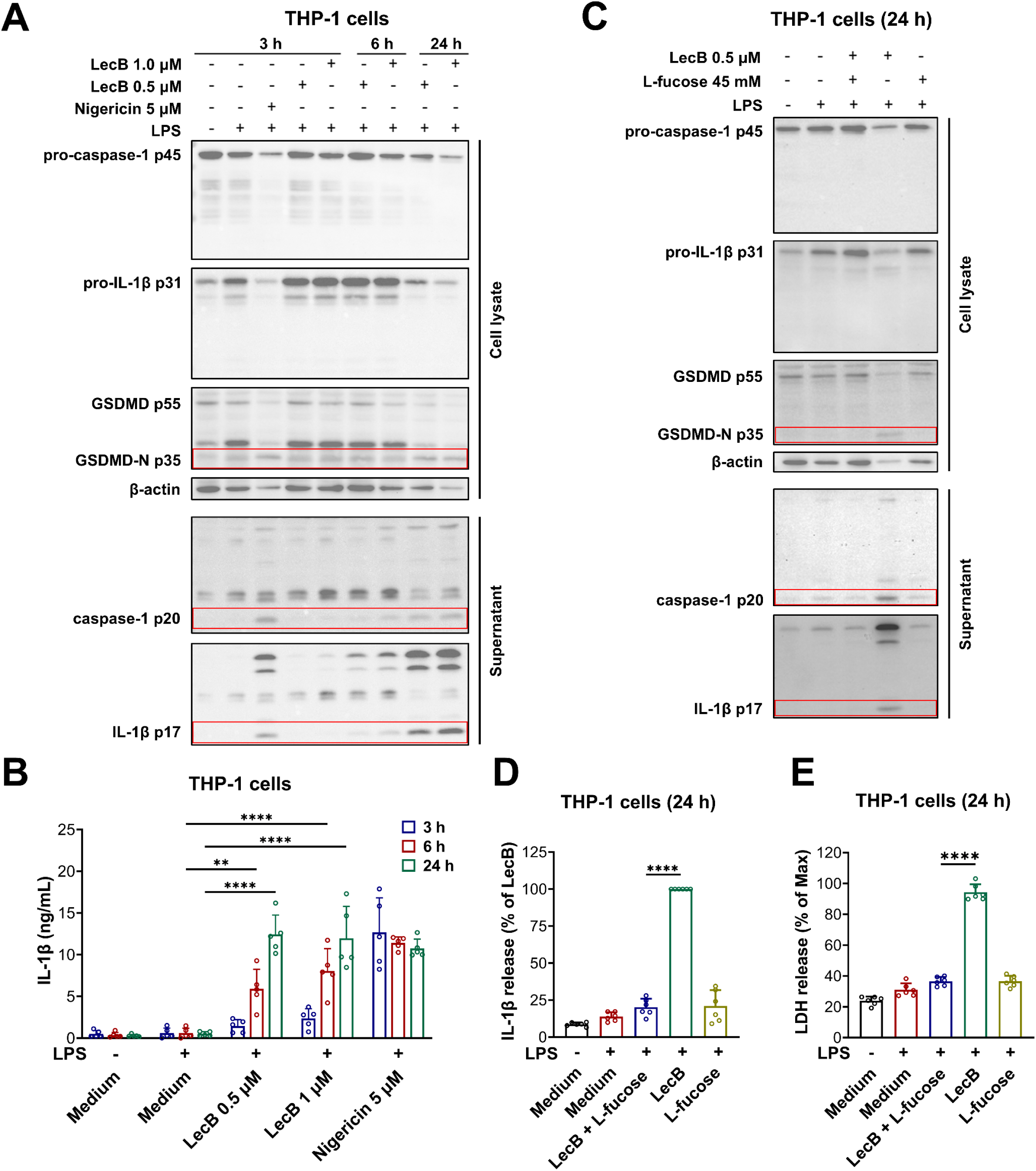
LecB triggers carbohydrate-dependent inflammasome activation in human macrophages. (A) Immunoblot of PMA-differentiated, unprimed or LPS-primed THP-1 cells stimulated with indicated concentrations of LecB for 3, 6 and 24 hours or Nigericin for 3 hours. (B) ELISA for IL-1β from supernatants of THP-1 cells stimulated with LecB or Nigericin for 3, 6 and 24 hours. (C–E) Unprimed or LPS-primed THP-1 cells were stimulated with LecB for 24 hours, with or without pre-incubation with L-fucose, or treated with L-fucose alone. Protein processing was analyzed by immunoblotting (C), IL-1β secretion by ELISA (D), and cell death by LDH release (E). Data information: Images in (A, C) are representative of n = 3 independent experiments. Graphs in (B) show mean ± SD from n = 5 independent experiments. Graphs in (D, E) show mean ± SD from n = 6 independent experiments. IL-1β secretion in (D) was normalized to the LecB-alone condition. Red boxes indicate cleavage products. *P* values for indicated comparisons were determined by two-way ANOVA in (B) and one-way ANOVA in (D, E), followed by Tukey’s multiple-comparisons test. \*\**p* < 0.01; \*\*\*\**p* < 0.0001.

Flow cytometry analysis demonstrated dose-dependent binding of both lectins to THP-1 cells. Despite having no effect on pyroptosis induction, LecA exhibited >15-fold stronger binding to THP-1 cells than LecB (Mean Fluorescence Intensity (MFI) 30,224 for LecA versus 1,686 for LecB at 0.5 μM (Figures S1C and S1D)). Binding of both lectins was reduced by their respective competing sugars (Figures S1E and S1F). Notably, LecB binding was almost completely abolished after pre-incubation with L-fucose (Figure S1F). Accordingly, presence of L-fucose blocked LecB-induced caspase-1 processing, prevented GSDMD cleavage, and reduced IL-1β maturation (Figure 1C). ELISA confirmed an approximately 80% reduction in IL-1β secretion when L-fucose was present, reaching levels comparable to the L-fucose-only controls (Figure 1D). Consistently, pyroptotic cell death was attenuated, with LDH release reduced by >60% (Figure 1E). Together, these data demonstrate that LecB induces inflammasome activation in THP-1 cells in a carbohydrate-binding-dependent manner and indicate that lectin binding strength alone does not predict inflammasome-activating potential.

### LecA triggers carbohydrate-dependent inflammasome activation in mouse macrophages

To explore whether *P. aeruginosa* lectins activate inflammasomes in other innate immune cells and across species, we next analyzed murine alveolar macrophage (AM)-like cells.^53^ In stark contrast to THP-1 cells (Figure 1), immunoblotting of cell lysates and supernatants showed that LecA induced a time- and dose-dependent inflammasome response, most pronounced at 24 hours, whereas LecB did not trigger detectable caspase-1 or IL-1β processing or GSDMD cleavage under any condition (Figures 2A and S2A). LecA-induced cleavage products were less prominent than those induced by Nigericin (Figure 2A). Consistently, ELISA demonstrated increasing IL-1β secretion over time in response to LecA, while LecB did not significantly elevate IL-1β as compared to LPS-primed controls (Figures 2B and S2B).

**Figure 2.**
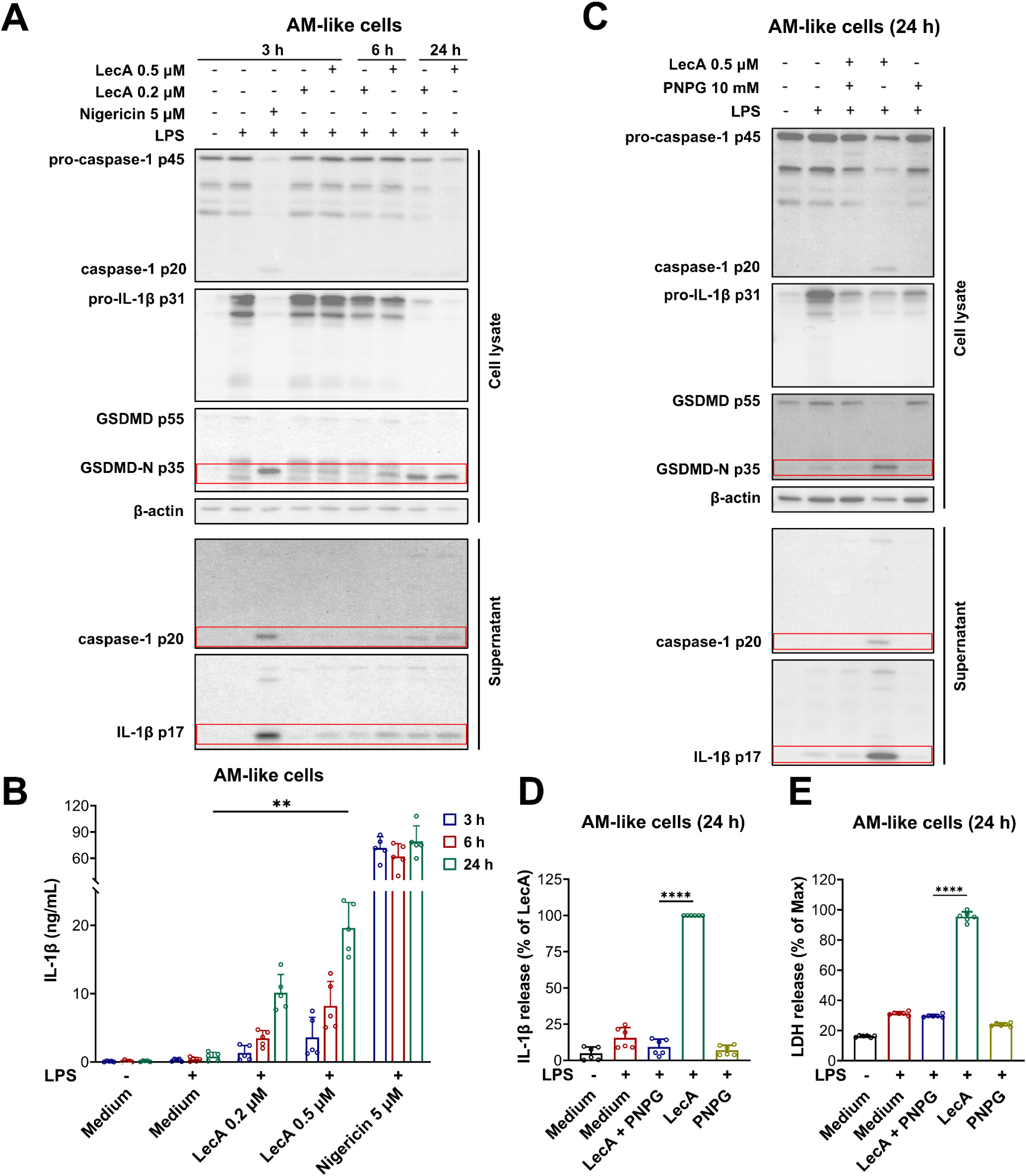
LecA triggers carbohydrate-dependent inflammasome activation in mouse macrophages. (A) Immunoblot of unprimed or LPS-primed AM-like cells stimulated with indicated concentrations of LecA for 3, 6 and 24 hours or Nigericin for 3 hours. (B) ELISA for IL-1β from supernatants of AM-like cells stimulated with LecA or Nigericin for 3, 6 and 24 hours. (C–E) Unprimed or LPS-primed AM-like cells were stimulated with LecA for 24 hours, with or without pre-incubation with PNPG, or treated with PNPG alone. Protein processing was analyzed by immunoblotting (C), IL-1β secretion by ELISA (D), and cell death by LDH release (E). Data information: Images in (A, C) are representative of n = 3 independent experiments. Graphs in (B) show mean ± SD from n = 5 independent experiments. Graphs in (D, E) show mean ± SD from n = 6 independent experiments. IL-1β secretion in (D) was normalized to the LecA-alone condition. *P* values for indicated comparisons were determined by two-way ANOVA in (B) and one-way ANOVA in (D, E), followed by Tukey’s multiple-comparisons test. \*\**p* < 0.01; \*\*\*\**p* < 0.0001.

Flow cytometry confirmed dose-dependent binding of both lectins to AM-like cells, with >790-fold stronger binding by LecA than LecB (MFI 122,815 for LecA versus 154 for LecB at 0.5 μM; Figures S2C and S2D). Consistent with D-galactose specificity, the competitive inhibitor p-nitrophenyl α-D-galactopyranoside (PNPG) reduced LecA binding by ∼35-fold (MFI 124,658 to 3,535), although residual binding remained above the medium-only control (MFI 53.7) (Figure S2E). In parallel, L-fucose further reduced the already weak LecB binding (MFI 159 to 81.9 (medium-only control MFI 53.5) (Figure S2F). Functionally, PNPG inhibited LecA-induced caspase-1 processing, IL-1β maturation and GSDMD cleavage (Figure 2C). It also reduced IL-1β secretion by ∼90% and LDH release from ∼95% to ∼30% (Figures 2D and 2E), indicating that similar to LecB-mediated inflammasome activation in THP-1 cells, LecA-driven inflammasome activation in AM-like cells depends on carbohydrate recognition.

Given that the glycosphingolipid Gb3 is a major LecA receptor,^21,54^ and we observed differing binding of LecA (Figure S1C and S2C), we hypothesized that Gb3 levels differ between murine AM-like cells and human THP-1 cells and performed targeted lipid profiling by LC-QqQ-MS. Although principal component analysis revealed pronounced global differences in total lipid content between cell types, accounting for ∼80% variance along PC1 which becomes apparent following clustering analysis of significantly altered lipids, total Gb3 levels were comparable. However, differences in other GlcCer-derived glycosphingolipids were observed between the two cell types, with higher levels of LacCer and Gb4 in THP-1 cells and Lc3 detected only in murine AM-like cells (Figures S3 and S4). The glucosylceramide (GlcCer) synthase inhibitor DL-threo-1-phenyl-2-palmitoylamino-3-morpholino-1-propanol (PPMP) was used to block the synthesis of GlcCer-based glycosphingolipids and thus to deplete cells of these molecules.^55,56^ Notably, PPMP treatment did not visibly reduce LecA-induced caspase-1 cleavage or release in AM-like cells (Figure S5), suggesting that inhibition of glycosphingolipid biosynthesis was insufficient to prevent LecA-induced inflammasome activation under these conditions.

We next investigated whether LecA-induced inflammasome activation could also be observed in other murine innate immune cells, including primary bone marrow-derived macrophages (BMDMs) and dendritic cells (BMDCs). In BMDMs, LecA induced a concentration-dependent increase in IL-1β secretion (Figure S6A). LDH release increased in parallel, slightly exceeding Nigericin-induced levels, although IL-1β secretion remained lower (Figure S6B). LecB had no detectable effect on IL-1β secretion and LDH release (Figures S6A and S6B). A similar pattern was observed in BMDCs: concentration-dependent LecA-induced IL-1β and LDH release, remaining below Nigericin positive control levels. LecB again showed only minimal activity (Figures S6C and S6D). Collectively, these data identify LecA as the dominant lectin driving glycan-dependent inflammasome activation in murine cells.

### Lectin binding triggers NLRP3 inflammasome activation and requires potassium efflux

Pyroptosis can be triggered by several inflammasomes. NLRP3 inflammasome has previously been implicated in *P. aeruginosa*-induced inflammatory responses.^48,57–59^ To determine whether the NLRP3 inflammasome is activated by *P. aeruginosa* lectins, we treated LPS-primed and lectin-stimulated THP-1 and AM-like cells with the selective NLRP3 inhibitor MCC950. MCC950 markedly suppressed lectin-induced caspase-1 and GSDMD cleavage and reduced IL-1β maturation (Figures 3A and 3B) and ELISA confirmed strong inhibition of IL-1β secretion (Figures 3C and 3D).

**Figure 3.**
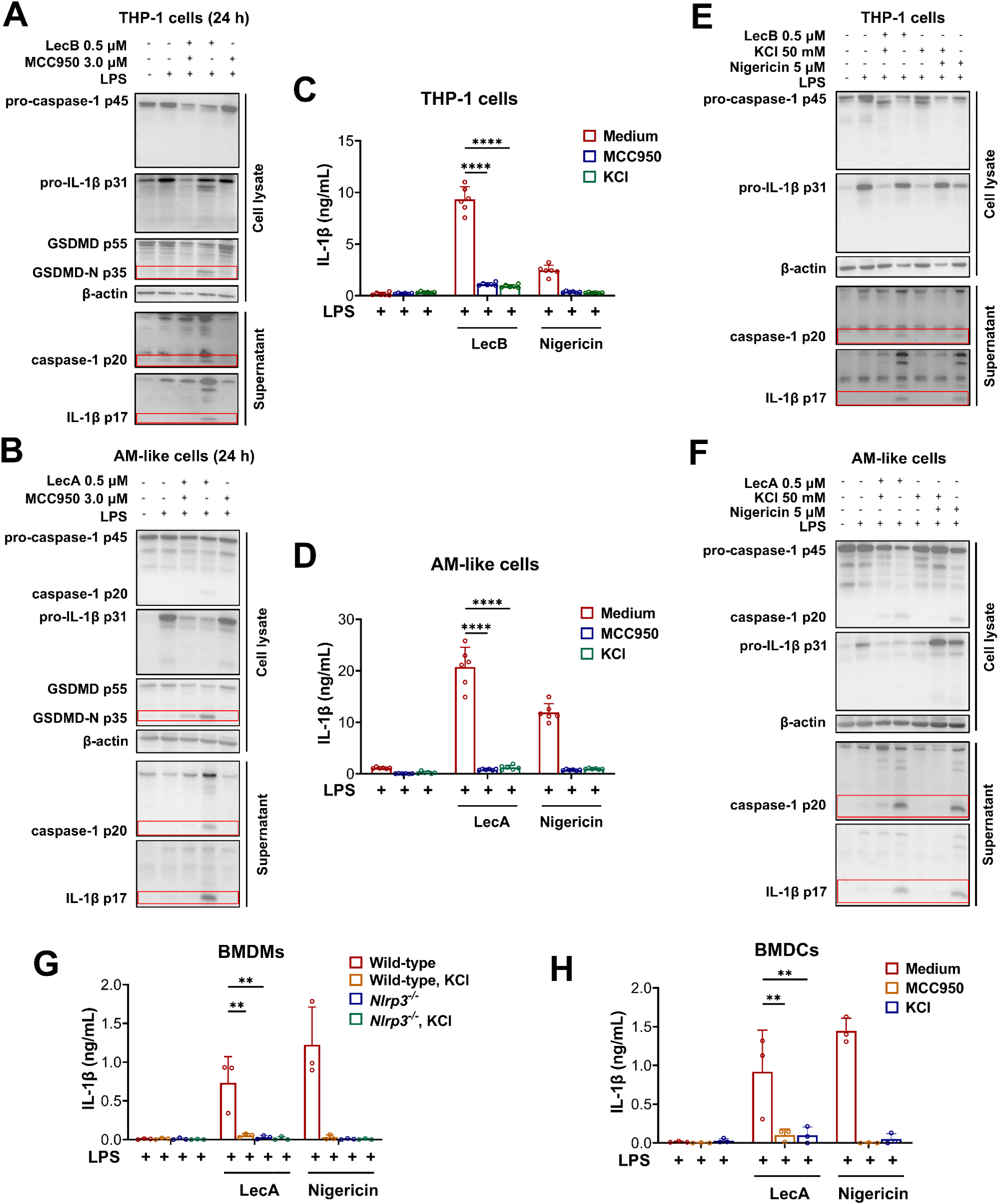
Lectin binding triggers NLRP3 inflammasome activation and requires potassium efflux. (A, B) Immunoblotting of PMA-differentiated THP-1 cells (A) and AM-like cells (B), unprimed or LPS-primed, stimulated with LecB or LecA, respectively, for 24 hours, with or without MCC950, or treated with MCC950 alone. (C, D) ELISA for IL-1β from supernatants of THP-1 cells (C) and AM-like cells (D) stimulated with LecB or LecA, respectively, for 24 hours, with or without MCC950 or KCl. Nigericin for 1 hour served as a positive control. (E, F) Immunoblot of THP-1 cells (E) and AM-like cells (F) treated as in (A, B), but with KCl instead of MCC950. Nigericin for 1 hour, with or without KCl, served as a positive control. (G) ELISA for IL-1β from supernatants of LPS-primed primary murine wild-type and *Nlrp3*^⁻/⁻^ BMDMs stimulated with LecA for 24 hours or Nigericin for 2 hours, with or without KCl, or treated with KCl alone. (H) ELISA for IL-1β from supernatants of LPS-primed primary murine BMDCs stimulated with LecA for 24 hours or Nigericin for 2 hours, with or without MCC950 or KCl, or treated with the respective inhibitor alone. Data information: Images in (A, B, E, F) are representative of n = 3 independent experiments. Graphs in (C, D) show mean ± SD from n = 6 independent experiments. Graphs in (G, H) show mean ± SD from n = 3 independent experiments. *P* values for indicated comparisons were determined by two-way ANOVA followed by Tukey’s multiple-comparisons test. \*\**p* < 0.01; \*\*\*\**p* < 0.0001.

Because potassium efflux is a key upstream trigger of NLRP3 activation, we tested its role in lectin responses.^60^ Increasing extracellular K⁺ prior to and during lectin-stimulation markedly reduced caspase-1 cleavage and IL-1β maturation and decreased IL-1β secretion (Figures 3C–3F), with inhibition comparable to MCC950.

To validate these findings in primary cells, including genetically in BMDMs, we generated primary wild-type (WT) and *Nlrp3*^⁻/⁻^ BMDMs and WT BMDCs and stimulated them with LecA for 24 hours. In BMDMs, IL-1β secretion was markedly reduced by absence of NLRP3, while in BMDCs it was strongly reduced by MCC950 treatment (Figures 3G and 3H). As expected, we observed a complete loss of IL-1β secretion by high extracellular K⁺ in WT BMDMs and BMDCs (Figure 3G and 3H). As control, Nigericin-induced IL-1β secretion was inhibited under the same conditions (Figure 3G and 3H). Together, these data demonstrate that *P. aeruginosa*-derived lectins induce macrophage and dendritic cell pyroptosis via the canonical potassium efflux-dependent NLRP3 pathway.

Reactive oxygen species (ROS), particularly from damaged mitochondria, also contribute to NLRP3 inflammasome activation.^61^ To probe this pharmacologically, we used the glutathione peroxidase-mimetic ROS scavenger Ebselen, which suppresses NLRP3-dependent IL-1β secretion in macrophages.^62^ In THP-1 and AM-like cells, Ebselen partially reduced lectin-induced caspase-1 cleavage and IL-1β maturation (Figures S7A and S7B) and decreased IL-1β secretion (Figures S7C and S7D). This effect was weaker than potassium efflux blockade (Figure 3C and 3D), indicating that ROS contribute but are not sufficient alone. In contrast, murine myeloid cells were more sensitive: Ebselen reduced LecA-induced IL-1β secretion to near control levels in BMDMs and to low residual levels in BMDCs (Figures S7E and S7F), suggesting cell type-dependent ROS involvement.

### Lectin binding induces ASC speck formation and NLRP3 relocalization

While not all inflammasomes require ASC or form a canonical ASC speck (CARD-containing sensors such as NLRC4 and murine NLRP1b can recruit and activate caspase-1 independently of ASC to induce pyroptosis),^63–65^ NLRP3 is ASC-dependent and its activation drives ASC speck formation.^66,67^ Therefore, we visualized ASC in LPS-primed THP-1 cells with immunofluorescence staining following stimulation with LecB for 24 hours. Confocal imaging and quantification showed robust ASC speck formation in ∼60% of cells, exceeding the response to Nigericin (∼30%; Figure 4A and 4B). Consistent with our biochemistry and ELISA data (Figure 1), this effect was markedly reduced by competitive blockade with L-fucose, which lowered speck formation to levels comparable to the L-fucose-only and the medium controls, indicating carbohydrate-dependent ASC oligomerization (Figures 4A and 4B). We next assessed LecA activity in murine AM-like cells. LecA induced pronounced ASC speck formation in ∼60% of cells, again exceeding Nigericin (∼45%). Consistent with glycan-dependent recognition and our biochemistry and ELISA data (Figure 2), PNPG efficiently blocked LecA-induced ASC speck formation, reducing it to control levels (Figures 4C and 4D). Similar to ASC, NLRP3 also shows subcellular relocalization upon inflammasome activation.^68^ Therefore, we examined NLRP3 localization by confocal microscopy following stimulation of LPS-primed THP-1 cells with fluorescently-labelled LecB. In control cells, NLRP3 displayed a diffuse cytosolic localization, followed by reduced signal intensity and sparse puncta formation over 6–24 hours following LecB stimulation (Figure S8). This relocalization was most evident at 24 hours, whereas LecA did not induce comparable changes in THP-1 cells (Figure S8). In murine AM-like cells, LecA induced a late relocalization of NLRP3, with perinuclear clustering observed only in a subset of cells at 24 hours, whereas NLRP3 remained diffusely distributed following LecB stimulation (Figure S9).

**Figure 4.**
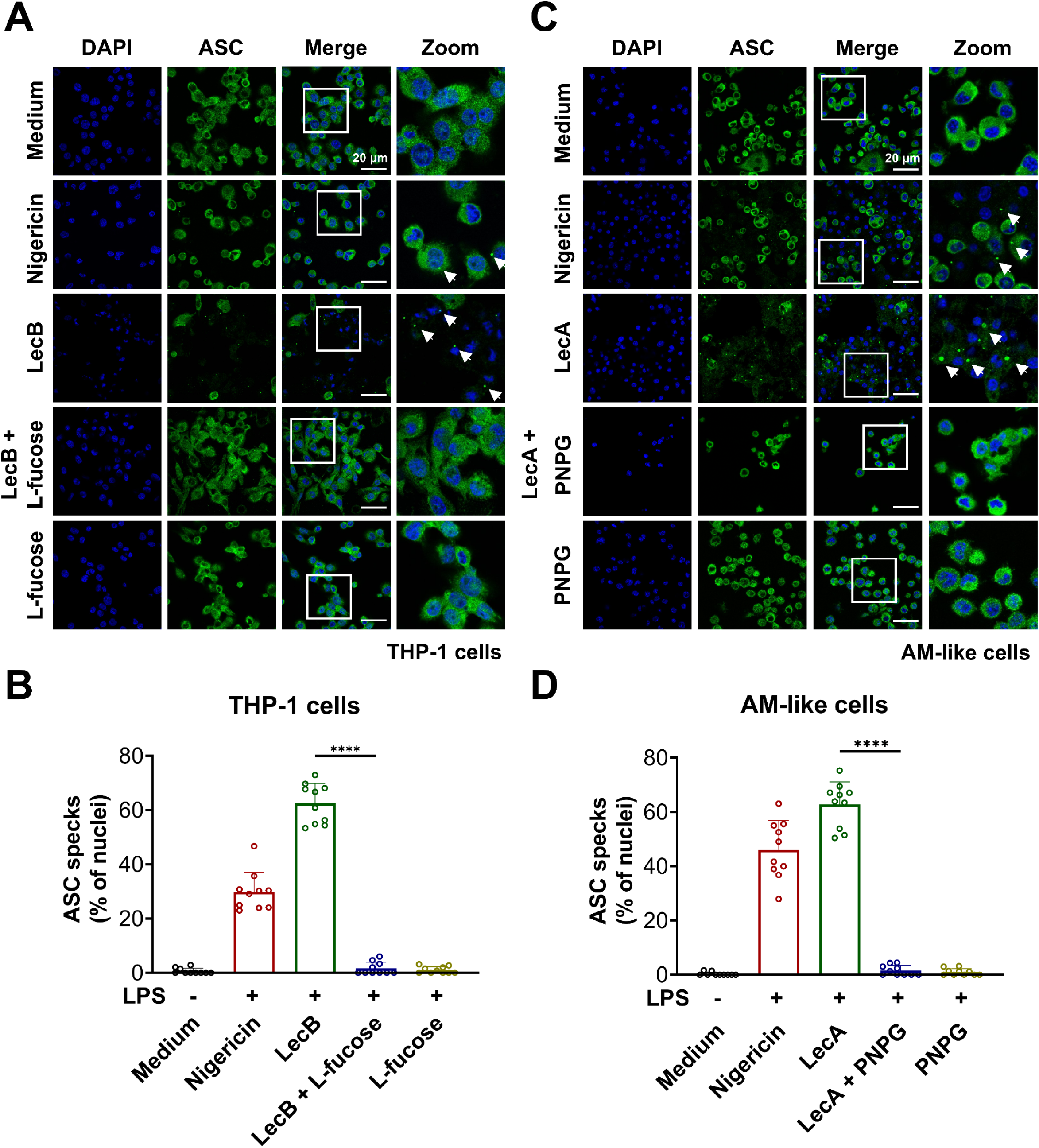
Lectin binding induces ASC speck formation. (A, C) Representative confocal images of PMA-differentiated, unprimed or LPS-primed THP-1 cells (A) and AM-like cells (C) stimulated with LecB or LecA, respectively, for 24 hours, with or without pre-incubation with the corresponding sugar inhibitor (L-fucose or PNPG). respectively. Nigericin for 1 hour served as a positive control. Cells were stained for ASC (green) and nuclei (DAPI, blue). Arrowheads indicate ASC specks. Scale bars, 20 μm. (B, D) Quantification of ASC speck formation in THP-1 cells (B) and AM-like cells (D) under the conditions shown in (A) and (C), respectively, expressed as ASC specks per nucleus (%). Each dot represents one field of view. Data information: Images in (A, C) are representative of n = 3 independent experiments. Data in (B, D) were pooled from n = 3 independent experiments, with a total of 10 randomly selected fields of view analyzed per condition, and are presented as mean ± SD. *P* values for indicated comparisons were determined by one-way ANOVA followed by Tukey’s multiple-comparisons test. \*\*\*\**p* < 0.0001.

Together, these data demonstrate that carbohydrate-dependent recognition of LecB and LecA in THP-1 cells and murine AM-like cells, respectively, drives ASC speck formation and NLRP3 spatial organization – morphological hallmarks of NLRP3 inflammasome activation.

### LecB drives carbohydrate-dependent activation of NLRP3 in PBMC-derived monocytes

To determine whether *P. aeruginosa* lectins can also activate the NLRP3 inflammasome in primary human immune cells, we stimulated LPS-primed peripheral blood mononuclear cell (PBMC)-derived monocytes with LecA or LecB for 24 hours. As previously observed using THP-1 cells (Figure 1), LecB induced robust IL-1β secretion, drastically exceeding the response to Nigericin, whereas LecA had no detectable effect (Figure 5A). LecB also caused lytic cell death as measured by LDH release, at levels comparable to Nigericin (Figure 5B), although lower than in THP-1 cells under similar conditions.

**Figure 5.**
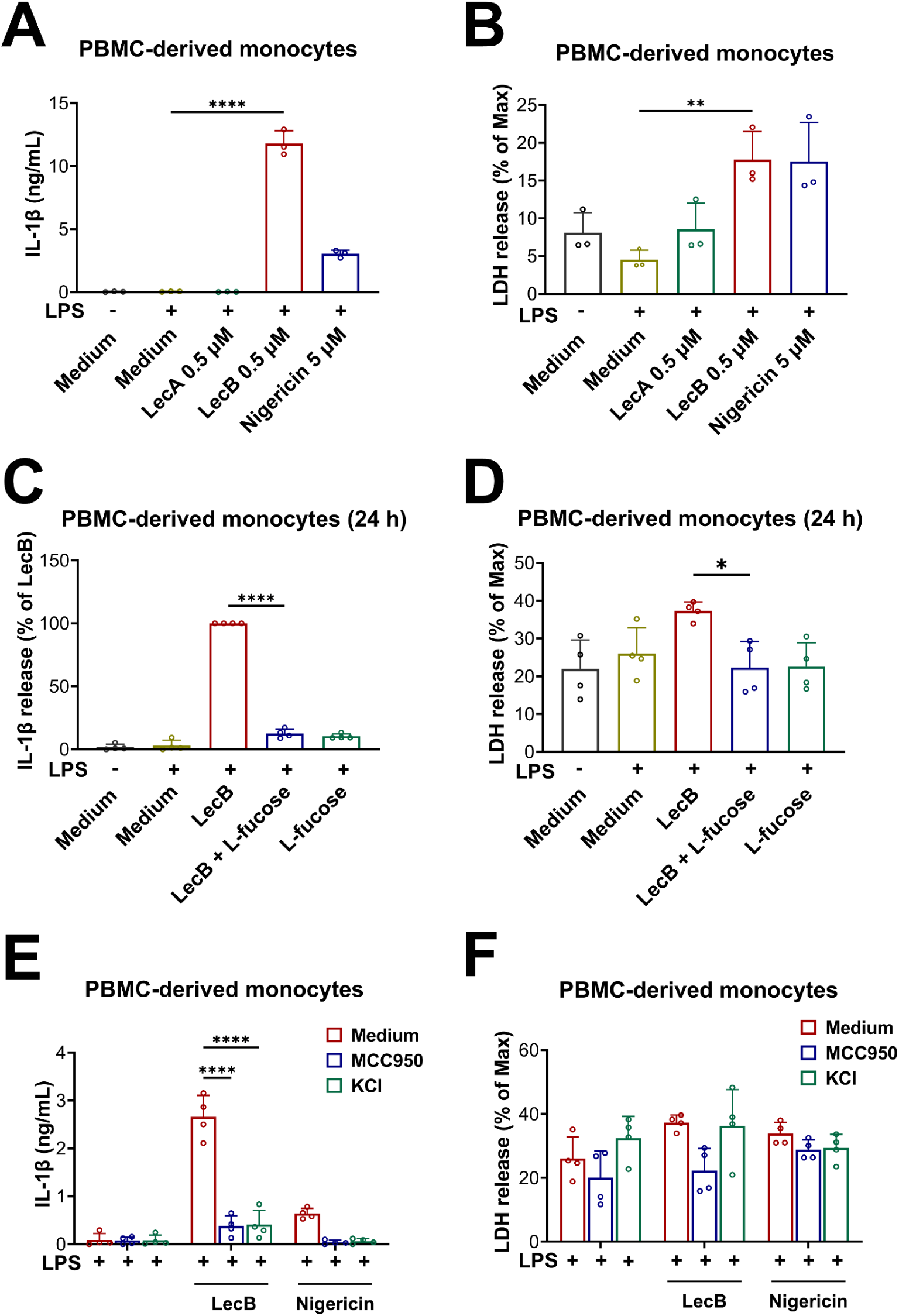
LecB drives carbohydrate-dependent activation of NLRP3 in PBMC-derived monocytes. (A, B) IL-1β secretion by ELISA (A) and cell death by LDH release (B) in unprimed or LPS-primed primary human PBMC-derived monocytes stimulated with LecA or LecB for 24 hours or Nigericin for 2 hours. (C, D) Cells were stimulated with LecB for 24 hours, with or without pre-incubation with L-fucose, or treated with L-fucose alone. IL-1β secretion (C) and cell death (D) were quantified by ELISA and LDH release, respectively. (E, F) LPS-primed cells were stimulated with LecB for 24 hours or Nigericin for 2 hours, with or without MCC950 or KCl, or treated with the respective inhibitor alone. IL-1β secretion (E) and cell death (F) were quantified by ELISA and LDH release, respectively. Data information: Graphs in (A, B) show mean ± SD from n = 3 donors. Graphs in (C–F) show mean ± SD from n = 4 donors. IL-1β secretion in (C) was normalized to the LecB-alone condition. *P* values for indicated comparisons were determined by one-way ANOVA in (A–D) and two-way ANOVA in (E, F), followed by Tukey’s multiple-comparisons test. \**p* < 0.05; \*\**p* < 0.01; \*\*\*\**p* < 0.0001.

To test carbohydrate dependence, monocytes from multiple donors were stimulated with LecB in the presence or absence of L-fucose which reduced LecB-induced IL-1β secretion by ∼90% close to the L-fucose-only control (Figure 5C). LDH release was similarly reduced from and comparable to L-fucose alone (Figure 5D). Inhibition of NLRP3 with MCC950 or blocking potassium efflux by elevated extracellular K⁺ strongly reduced LecB-induced IL-1β secretion to near LPS-controls, consistent with potassium efflux-dependent NLRP3 activation observed before (Figure 5E). As control, Nigericin responses were similarly inhibited (Figure 5E). In contrast, LDH release was only modestly affected by MCC950 and largely insensitive to high extracellular K⁺ after LecB stimulation, while Nigericin-induced LDH release was only slightly reduced (Figure 5F).

Collectively, these data indicate that LecB-driven inflammasome responses in primary human monocytes require lectin-glycan interactions, whereas the accompanying lytic cytotoxicity is only partially NLRP3-dependent and likely engages additional inflammasome-independent death pathways.

### Integrin α5β1 contributes to LecB-triggered NLRP3 inflammasome activation in THP-1 cells

To identify host surface glycoproteins involved in LecB-triggered inflammasome activation, we examined the association of LecB with integrin β1 (Figure 6A), which had previously been identified as an interaction partner in H1299 lung epithelial cells.^32,34^ THP-1 cells were incubated with biotinylated LecB (LecB-biotin), lysed, and LecB-containing complexes were isolated by streptavidin pull-down. Immunoblot analysis of input lysate, unbound flow-through, and pull-down fractions showed integrin β1 in the lysate and remaining in the flow-through, while a distinct fraction was specifically recovered in the pull-down in the presence of LecB-biotin (Figure 6A). No integrin β1 signal was detected in pull-downs controls. Given the fucose-binding specificity of LecB, we next assessed whether integrin β1 glycosylation contributes to this interaction. THP-1 lysates with or without PNGase F treatment were analyzed by immunoblotting for integrin β1 and by an overlay assay using fluorescent LecB (LecB-AF488). PNGase F treatment induced a clear downward mobility shift of integrin β1, consistent with removal of N-linked glycans (Figure 6B). In untreated lysates, LecB-AF488 showed strong binding, including a prominent signal co-migrating with glycosylated integrin β1. This signal was lost after PNGase F treatment, and overall LecB binding was markedly reduced (Figure 6B). These data indicate that, as previously observed in epithelial cells, LecB also associates with integrin β1 on the surface of THP-1 cells, consistent with glycan-dependent recognition at the integrin β1.

**Figure 6.**
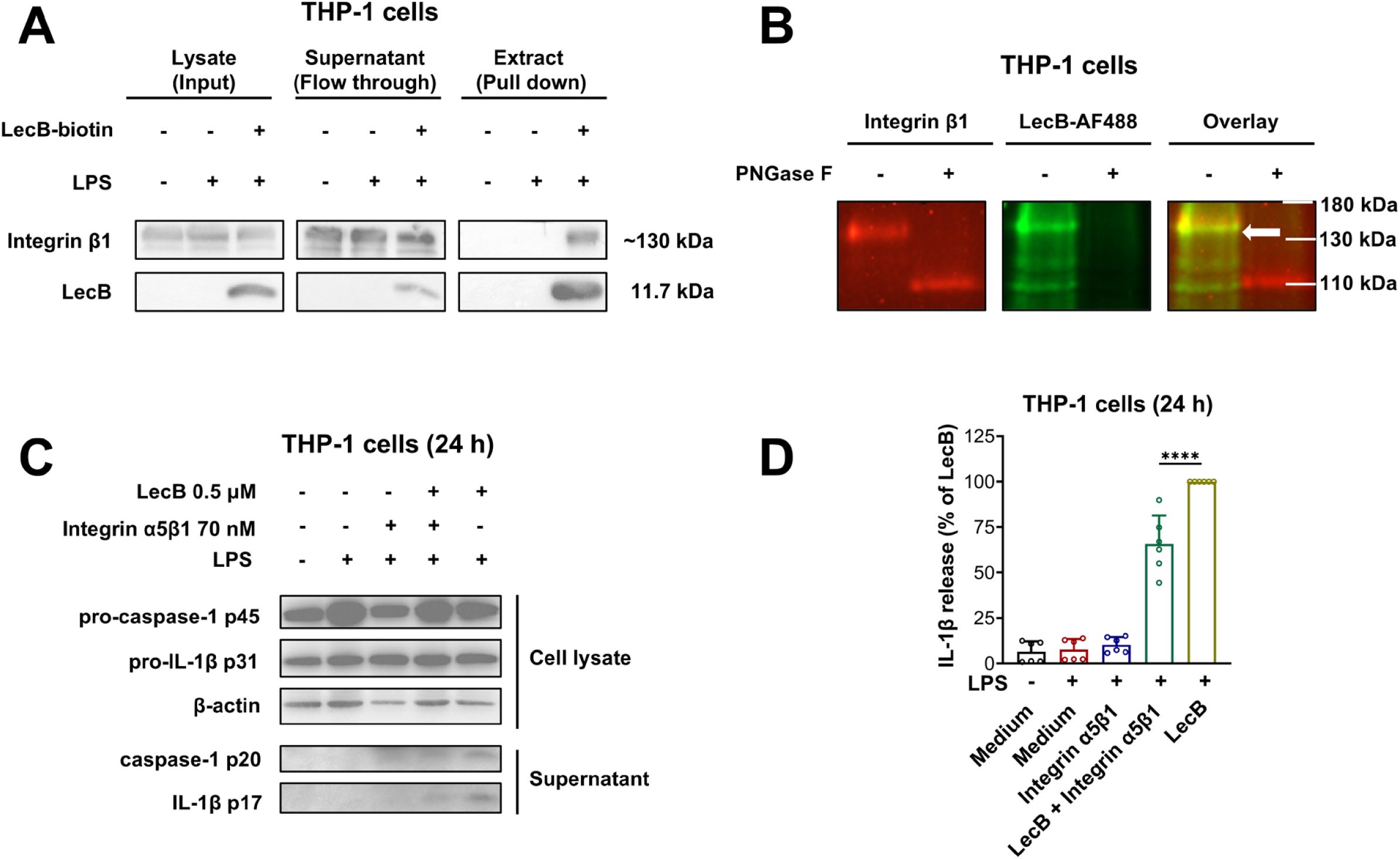
Integrin α5β1 contributes to LecB-triggered NLRP3 inflammasome activation in THP-1 cells. (A) Streptavidin pull-down of LecB-biotin-containing complexes from PMA-differentiated, unprimed or LPS-primed THP-1 cells, followed by immunoblotting for integrin β1 and LecB. Input, flow-through and pull-down fractions are shown. (B) Immunoblotting for integrin β1 and a LecB-AF488 overlay assay using THP-1 cell lysates with or without PNGase F treatment. White arrows indicate LecB-AF488 signal co-migrating with integrin β1. (C, D) Unprimed or LPS-primed THP-1 cells were stimulated with LecB for 24 hours, with or without anti-integrin α5β1 antibody. Protein processing was analyzed by immunoblotting (C) and IL-1β secretion by ELISA (D). Data information: Images in (A–C) are representative of n = 3 independent experiments. Data in (D) are presented as mean ± SD from n = 6 independent experiments. IL-1β secretion in (D) was normalized to the LecB-alone condition. *P* values for indicated comparisons in (D) were determined by one-way ANOVA followed by Tukey’s multiple-comparisons test. \*\*\*\**p* < 0.0001.

To test functional relevance for the induction of pyroptosis, a blocking anti-integrin α5β1 antibody was applied during LecB stimulation. Antibody treatment reduced caspase-1 processing and IL-1β maturation (Figure 6C), and ELISA confirmed decreased IL-1β secretion, although levels remained above the LPS-primed control (Figure 6D). In contrast, an RGD peptide, which competes for the canonical integrin ligand-binding site, did not inhibit LecB-induced inflammasome activation even at high concentration (Figure S10), suggesting that LecB engages α5β1 independently of the RGD-binding interface. Together, these data indicate a role for α5β1 integrin and N-linked glycans in LecB-driven inflammasome activation.

### *P. aeruginosa*-induced inflammasome activation in THP-1 cells is attenuated by L-fucose

To assess physiological relevance and whether lectins contribute to *P. aeruginosa* induced macrophage pyroptosis, we infected PMA-differentiated, LPS-primed THP-1 cells with live *P. aeruginosa* PAO1 (MOI 0.1 or 1) for 6 or 24 hours. At 6 hours, IL-1β secretion increased, most prominently at MOI 1, while caspase-1 processing was not detected (Figure 7A). This early IL-1β release may involve alternative processing mechanisms, such as direct cleavage of pro-IL-1β by the *P. aeruginosa* metalloprotease LasB.^69^ By 24 hours, infection induced clear caspase-1 processing together with IL-1β secretion; caspase-1 cleavage was more pronounced at MOI 0.1, whereas IL-1β levels were higher at MOI 1 (Figure 7B). The results obtained at MOI 1 after 24 hours should be interpreted with caution due to the absence of a detectable β-actin signal. Nevertheless, LecB-induced inflammasome activation was clearly observed at MOI 0.1. Co-incubation with L-fucose partially reduced caspase-1 processing and IL-1β release at 24 hours (Figure 7B), consistent with a contribution of fucose-binding factors such as LecB. To determine whether ectopic expression of LecB is sufficient to trigger inflammasome activation in a bacterial context, THP-1 cells were infected with the LecB-expressing *E. coli* BL21(DE3) strain (MOI 100), which induced caspase-1 processing and IL-1β release at 24 hours (Figure 7C). These findings support a functional role for LecB in inflammasome activation during *P. aeruginosa* infection.

**Figure 7.**
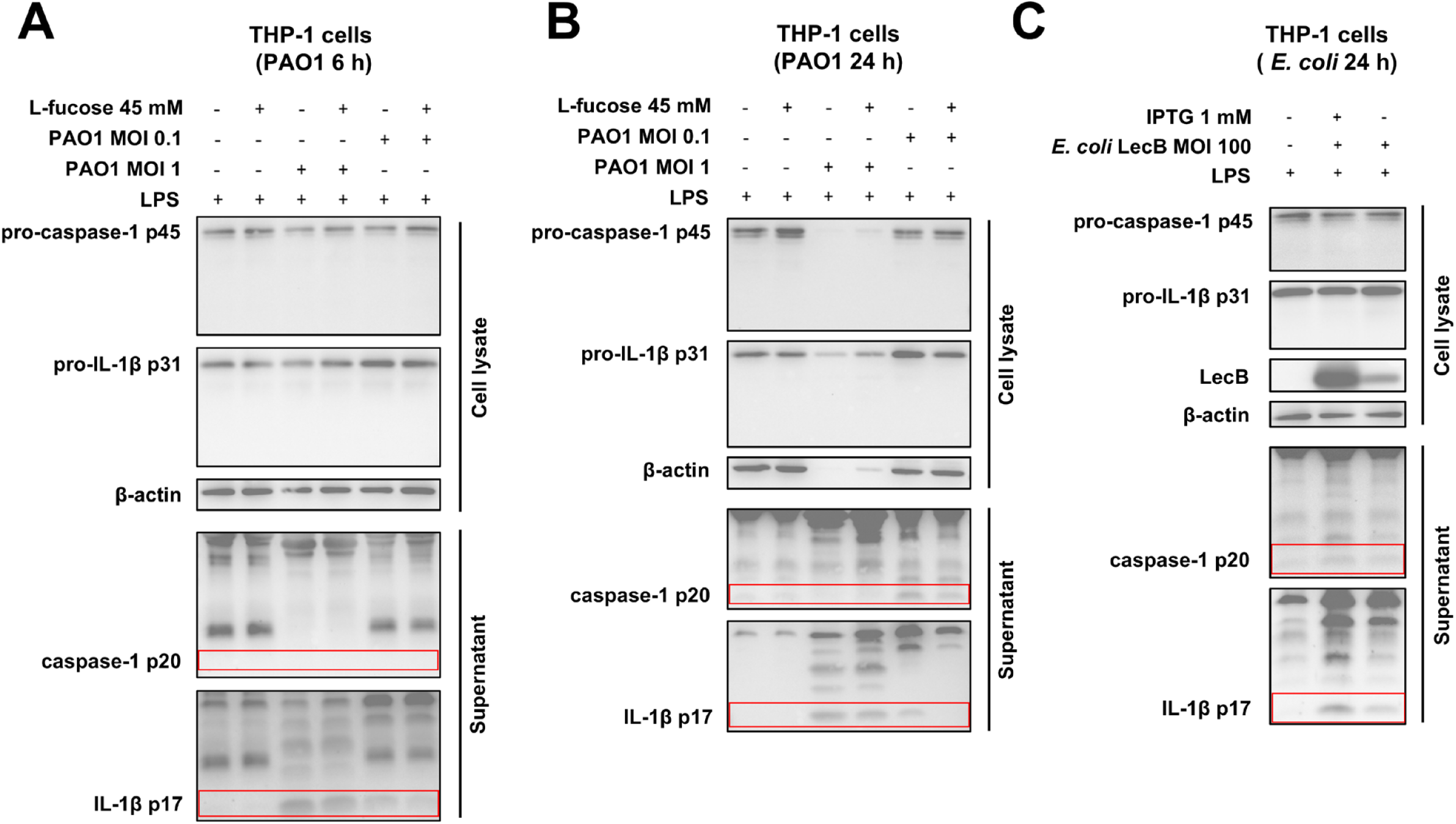
*P. aeruginosa*-induced inflammasome activation in THP-1 cells is attenuated byL-fucose. (A, B) Immunoblot of PMA-differentiated, LPS-primed THP-1 cells infected with live *P. aeruginosa* PAO1 at the indicated MOI for 6 hours (A) or 24 hours (B), with or without pre-incubation of PAO1 with L-fucose. (C) Immunoblot of LPS-primed THP-1 cells infected for 24 hours with LecB-expressing *E. coli* BL21(DE3) cultured with or without IPTG induction. Data information: Images in (A–C) are representative of n = 3 independent experiments.

## DISCUSSION

As a major opportunistic pathogen associated with multidrug-resistant infections, *Pseudomonas aeruginosa* exploits a sophisticated repertoire of virulence factors to subvert host immunity and cause life-threatening infections in immunocompromised patients.^4,70–72^ Central to this immunomodulatory strategy is a striking functional dichotomy, where different virulence factors exert opposite effects on host inflammasome pathways. For instance, while secreted effector proteins like ExoU and ExoS actively suppress intracellular signaling, structural components of the bacterium, such as flagellin and T3SS needle/rod proteins, trigger NLRC4 inflammasome activation via NAIP recognition.^73–76^ This intricate interplay underscores a finely tuned evolutionary arms race between bacterial evasion and host detection. However, host defense is far from passive. Host cells have co-evolved sophisticated cell-autonomous and intercellular strategies to counter these bacterial evasive maneuvers,^77,78^ actively converting the pathogeńs own virulence factors into trigger for immunological surveillance.^79^

Here, we identify the *P. aeruginosa* lectins LecA and LecB, traditionally recognized for driving biofilm formation, epithelial barrier disruption, and bacterial internalization,^21,24,34,80–82^ as primary targets of this host surveillance. We demonstrate that innate immune cells exploit cell-surface glycans to capture LecA and LecB, effectively turning these carbohydrate-binding proteins into a molecular trigger that mediates NLRP3 inflammasome activation. This finding reveals a novel paradigm in host-pathogen dynamics: The human immune system not only senses foreign microbial carbohydrate structures, but also strategically deploys its own carbohydrates as bait to intercept bacterial lectins, transforming a bacterial virulence factor into an autonomous platform for immune activation. While LecB elicits stronger immune responses in human THP-1 cells and PBMC-derived monocytes, LecA preferentially activates the NLRP3 inflammasome in murine AM-like cells, BMDMs, and BMDCs. This distinct functional divergence may partly reflect differences in cell-surface glycan repertoires and lectin receptor availability among the human and murine cell types examined.^83–85^

Based on our time-course experiments, lectin-induced inflammasome activation required prolonged stimulation and became prominent at 24 hours, in contrast to conventional rapid NLRP3 stimuli.^86,87^ These delayed kinetics suggest that sustained lectin-host cell interactions and/or distinct intracellular trafficking routes may be required to cross the activation threshold for NLRP3 signaling. Importantly, the delayed kinetics may allow substantial IL-1β production before terminal cell death. This possibility is reminiscent of oxidized-phospholipid-induced hyperactivation in dendritic cells, in which inflammasome-dependent IL-1β release can occur without immediate pyroptosis and support subsequent adaptive immune responses.^88^ A defining feature of this response is its strict dependence on carbohydrate recognition. Inhibiting LecB or LecA binding markedly reduced inflammasome activation, indicating that lectin-glycan interactions function not simply as adhesive contacts but as an upstream determinant of immune activation. This concept parallels the action of other glycan-binding bacterial toxins. For instance, Shiga toxin activates the NLRP3 inflammasome via engagement with its specific receptor Gb3^89^, while the cholera toxin B subunit (CTB), which targets the ganglioside GM1, drives IL-1β production through both the pyrin and NLRP3 inflammasomes.^90^ Collectively, our results support a broader paradigm in which the host immune system exploits microbial glycan-binding factors as a primary signal to license inflammasome assembly.

Mechanistically, our data demonstrate that *P. aeruginosa* lectins activate the canonical NLRP3 inflammasome through a potassium efflux-dependent pathway. Furthermore, our results implicate glycosylated β1-integrin complexes as important upstream mediators of LecB-driven NLRP3 inflammasome activation. These observations align closely with prior work showing that LecB targets glycosylated β1-integrin during host cell interactions and internalization,^34^ as well as evidence linking α5β1-integrin to NLRP3 activation in response to the *Treponema denticola* surface protein Td92 in human macrophages.^91^ Such parallels suggest that distinct microbial surface factors may converge on integrin-dependent pathways to promote inflammasome activation. A potential mechanistic link comes from the ability of LecB to modulate Piezo1, a mechanosensitive, nonselective cation channel, in a time- and carbohydrate-dependent manner.^92^ Given that Piezo1 gating profoundly potentiates the NLRP3 inflammasome during *Candida albicans* infection,^93^ LecB-induced alterations in Piezo1 activity may provide an additional regulatory input. We hypothesize that this mechanical and biochemical crosstalk licenses NLRP3 activation by driving sustained Ca²⁺ influx and subsequent secondary ionic disturbances, ultimately triggering the threshold-level potassium efflux required for mature inflammasome assembly.

In contrast, LecA-induced membrane remodeling^94^ may influence mechanosensitive ion-channel activity. Indeed, LecA has been shown to enhance the activity of the mechanosensitive potassium channel TREK-1 in response to mechanical and electrical stimuli, without engaging Piezo1.^95^ Given the recently identified role of TREK-1 in NLRP3 activation in murine macrophages,^96^ these findings suggest a potential link between LecA-mediated membrane perturbations, potassium channel regulation, and NLRP3 inflammasome activation.

Collectively, our findings support a paradigm wherein LecB and LecA are trapped in distinct, glycan-dependent host topologies to converge on canonical NLRP3 inflammasome activation. Dissecting the precise proximal signaling events that bridge these discrete lectin-receptor interactions to uniform downstream ion fluxes remains an essential objective for future research. Beyond this shared potassium efflux-dependent response, the differential sensitivity to Ebselen suggests that oxidative stress may contribute differently to lectin-induced inflammasome activation across cell types. In this regard, recent studies have demonstrated that mitochondrial electron transport chain (ETC) activity can license NLRP3 activation via phosphocreatine (PCr)-dependent ATP production, completely independent of ROS generation.^97^ Whether a similar metabolic mechanism contributes to lectin-induced inflammasome activation remains to be determined.

To assess the relevance of these glycan-dependent mechanisms during infection, we examined the host response to live *P. aeruginosa* (strain PAO1). The delayed inflammasome activation observed during infection resembled the kinetics induced by purified LecB, supporting the physiological relevance of lectin-mediated host recognition. Although L-fucose partially reduced inflammasome activation during *P. aeruginosa* infection, the multifaceted virulence repertoire of this pathogen complicates direct attribution to LecB alone.

In conclusion, this study identifies *P. aeruginosa* lectins as key drivers of glycan-mediated NLRP3 inflammasome activation. Species-specific differences in glycosylation profiles likely explain the distinct responses observed between human and murine cells,^98,99^ emphasizing that caution must be exercised when extrapolating murine in vivo data to human immune contexts.^100,101^

Ultimately, elucidating this glycan-dependent axis not only advances our fundamental understanding of host-pathogen interactions but also highlights bacterial lectins as high-value therapeutic targets for the modulation of aberrant inflammation. Future investigations should focus on identifying the specific host glycan receptors involved and delineating the upstream signaling pathways that bridge lectin engagement to NLRP3 activation.

## ACKNOWLEDGMENTS

This study was supported by the German Research Foundation (DFG) under Germany’s Excellence Strategy (CIBSS – EXC-2189 – Project ID 390939984) and the DFG grant Major Research Instrumentation (438033605), as well as the Ministry of Science, Research and the Arts of the State of Baden-Württemberg (Az: 33–7532.20). We would like to thank Prof. Marina Freudenberg for generously providing murine alveolar macrophage (AM)-like cells. SZ, XW and SS acknowledge support from the China Scholarship Council.

## AUTHOR CONTRIBUTIONS

Conception and design of the research: SZ, OM, WR; funding acquisition: BK, OG, WR; investigation: SZ, OM, XW, SW, SL; methodology: SZ, OM, XW, SW, SL, SS, CD, NH, LW, BK, OG, DF, WR; supervision: OM, BK, OG, DF, WR; writing - original draft: SZ, OM, WR; writing - review and editing: SZ, OM, SW, XW, SL, SS, CD, NH, LW, BK, OG, DF, WR. All authors agreed on the final version of the manuscript.

## STAR★METHODS

## KEY RESOURCES TABLE

| REAGENT or RESOURCE | SOURCE | IDENTIFIER |
|---|---|---|
| **Antibodies** |  |  |
| anti-Caspase-1 (p20) (human) mAb (Bally-1) | Adipogen | Cat# AG-20B-0048; RRID: AB_2490257 |
| anti-Caspase-1 (p20) (mouse) mAb (Casper-1) | Adipogen | Cat# AG-20B-0042, RRID: AB_2490248 |
| anti-NLRP3/NALP3 mAb (Cryo-2) | Adipogen | Cat# AG-20B-0014-C100, RRID: AB_2490202 |
| IL-1β (D3U3E) | Cell Signaling Technology | Cat# 12703, RRID: AB_2737350 |
| Mouse IL-1 beta/IL-1F2 Antibody | R&D Systems | Cat# AF-401-NA, RRID: AB_416684 |
| anti-GSDMD (126-138) antibody produced in rabbit | Sigma-Aldrich | Cat# G7422, RRID: AB_1850381) |
| anti-GSDMD antibody [EPR19828] | Abcam | Cat# ab209845, RRID: AB_2783550) |
| Mouse anti-integrin β1 | Millipore | Cat# MAB2259Z, RRID: AB_94616 |
| Anti-Integrin α5β1, clone JBS5 | Sigma-Aldrich | Cat# MAB1969, RRID: AB_94472 |
| anti-beta-Actin | Sigma-Aldrich | Cat# A5316, RRID: AB_476743 |
| anti-Asc pAb (AL177) (ATTO 647N) | Adipogen | Cat# AG-25B-0006TS, RRID: AB_2490442 |
| anti-LecB | Eurogentec | N/A |
| Goat anti-Rabbit IgG (H+L), Alexa Fluor™ 647 | Thermo Fisher Scientific | Cat# A-21245, RRID: AB_2535813 |
| Goat anti-Mouse IgG (H+L), Alexa Fluor™ 647 | Thermo Fisher Scientific | Cat# A-21236, RRID: AB_2535805 |
| anti-mouse IgG, HRP-linked Antibody | Cell Signaling Technology | Cat# 7076, RRID: AB_330924 |
| anti-rabbit IgG, HRP-linked Antibody | Cell Signaling Technology | Cat# 7074, RRID: AB_2099233 |
| **Bacterial strains** |  |  |
| *Pseudomonas aeruginosa* PAO1 | Gift from Sophie de Bentzmann, Marseille | N/A |
| *Escherichia coli* LecA | Gift from Anne Imberty, Grenoble | N/A |
| *Escherichia coli* LecB | Gift from Anne Imberty, Grenoble | N/A |
| **Chemicals, proteins, etc.** |  |  |
| RPMI 1640 w/o L-Glutamine | anprotec | Cat# AC-LM-0060 |
| RPMI 1640 Medium, GlutaMAX™ Supplement | Life Technologies GmbH | Cat# 61870010 |
| Fetal bovine serum | Gibco | Cat# P30-3302 |
| Fetal calf serum | Life Technologies GmbH | Batch no. B2695491RP (no catalog number) |
| L-Glutamine | Capricorn Scientific | Cat# GLN-B |
| Penicillin-streptomycin | anprotec | Cat# AC-AB-0042 |
| HEPES | gibco | Cat# 15630080 |
| EDTA | Carl Roth | Cat# 8040.1 |
| DPBS | anprotec | Cat# AC-BS-0002 |
| Ficoll-Paque™ PLUS density gradient media | VWR International GmbH | Cat# 17144002 |
| Recombinant Mouse GM-CSF (carrier-free) | BioLegend | Cat# 576304 |
| Recombinant murine GM-CSF | Immunotools | Cat# 12343128 |
| Recombinant human M-CSF | 7BioScience GmbH | Cat# 7B-hMCSF_1MG |
| PMA | Sigma-Aldrich | Cat# P1585 |
| E. coli K12 ultra-pure LPS | InvivoGen | Cat# tlrl-peklps |
| Lipopolysaccharides from Escherichia coli O111:B4 | Sigma-Aldrich | Cat# L4391 |
| Nigericin | Sigma-Aldrich | Cat# N7143 |
| MCC950 | Adipogen | Cat# AG-CR1-3615 |
| KCl | Sigma-Aldrich | Cat# P9333 |
| KCl | Carl Roth | Cat# 6781.1 |
| Ebselen | Sigma-Aldrich | Cat# E3520 |
| Ebselen | Biomol GmbH | Cat# Cay70530 |
| PNGase F | Sigma-Aldrich | Cat# P7367 |
| LI-COR Protein-Free Blocking Buffer | LICORBIO | Cat# 927-80001 |
| RGD peptide (GRGDNP) | MedChemExpress | Cat# HY-P1740A |
| Pierce™ magnetische Streptavidin-Beads | Thermo Fisher Scientific | Cat# 88817 |
| L-Fucose | Sigma-Aldrich | Cat# F2252 |
| p-Nitrophenyl α-D-galactopyranoside (PNPG) | Sigma-Aldrich | Cat# N0877 |
| DL-threo-1-Phenyl-2-palmitoylamino-3-morpholino-1-propanol (PPMP) | Santa Cruz | Cat# sc-205655 |
| Pierce™ D-Galactose Agarose | Thermo Fisher Scientific | Cat# 20372 |
| D-Mannose Agarose | Sigma-Aldrich | Cat# M6400 |
| DMSO | Carl Roth | Cat# A994.2 |
| SDS ultra pure | Carl Roth | Cat# 2326.2 |
| DL-Dithiothreitol (DTT) | Sigma-Aldrich | Cat# 43819 |
| TRIS | Carl Roth | Cat# 4855.5 |
| ROTIPHORESE®NF-Acrylamide/Bis-solution 30 (29:1) | Carl Roth | Cat# A124.1 |
| TEMED | Carl Roth | Cat# 2367.1 |
| PageRuler™ Prestained Protein Ladder, 10 to 180 kDa | Thermo Fisher Scientific | Cat# 26617 |
| Ponceau S | Sigma-Aldrich | Cat# P3504 |
| ROTI®Fair 10x PBS 7.4 | Carl Roth | Cat# 1105.1 |
| Tween 20 | Carl Roth | Cat# 9127.1 |
| Powdered milk | Carl Roth | Cat# T145.3 |
| Sodium hydrogen carbonate | Carl Roth | Cat# HN01.1 |
| Bovine Serum Albumin | Sigma-Aldrich | Cat# A9647 |
| Saponin | Sigma-Aldrich | Cat# 84510 |
| Ammonium chloride | Carl Roth | Cat# K298.1 |
| Pierce™ 16% Formaldehyde (w/v), Methanol-free | Thermo Fisher Scientific | Cat# 28908 |
| DMSO, Anhydrous | Thermo Fisher Scientific | Cat# D12345 |
| DAPI | Carl Roth | Cat# 6335 |
| Alexa Fluor™ 488 NHS-Ester (Succinimidylester) | Thermo Fisher Scientific | Cat# A20000 |
| IPTG | Carl Roth | Cat# 2316.4 |
| LB Broth | Carl Roth | Cat# 6669.2 |
| Ampicillin | Carl Roth | Cat# K029.5 |
| 2-Propanol | VWR | Cat# 84881.320 |
| Acetonitrile | VWR | Cat# 83640.400 |
| Formic acid | VWR | Cat# 84865.290 |
| Chloroform | VWR | Cat# 83627.320 |
| Ammonium formate | VWR | Cat# 84884.260 |
| Methanol | VWR | Cat# 85800.320 |
| Shiga Toxin 1, B subunit | Sigma-Aldrich | Cat# SML0562 |
| LecA | This study | N/A |
| LecB | This study | N/A |
| **Critical commercial assays** |  |  |
| Mouse IL-1 beta Uncoated ELISA Kit | Thermo Fisher Scientific | Cat# 88-7013A-76 |
| Human IL-1 beta Uncoated ELISA Kit | Thermo Fisher Scientific | Cat# 88-7261-76 |
| cytotox 96® non-radioactive cytotoxicity assay | Promega | Cat# G1780 |
| Thermo Scientific™ EZ-Link™ NHS-PEG4 Biotinylation Kit | Thermo Fisher Scientific | Cat#10701185 |
| Novex™ ECL Chemiluminescent Substrate Reagent Kit | Thermo Fisher Scientific | Cat# WP20005 |
| Westar Supernova | Cyangen | Cat# XLS3,0100 |
| **Experimental models: Cell lines** |  |  |
| THP-1 cells | ATCC | N/A |
| Murine alveolar macrophage (AM)-like cells | Gift from Marina Freudenberg, Max Planck Institute, Freiburg, Germay | N/A |
| Murine bone marrow-derived dendritic cells (BMDCs) | This study | N/A |
| Murine bone marrow-derived macrophages (BMDMs) | This study | N/A |
| **Mice** |  |  |
| Mice, C57BL/6 background, wild-type | CEMT, Freiburg | N/A |
| Mice, C57BL/6 background, *Nlrp3*<sup>-/-</sup> | J. Tschopp, Lausanne; Martinon et al. | https://doi.org/10.1038/nature04516 |
| **Biological samples** |  |  |
| Human blood samples | This study | N/A |
| **Software and algorithms** |  |  |
| GraphPad Prism | GraphPad Software | N/A |
| FlowJo | BD Biosciences | N/A |
| ImageJ Fiji | https://fiji.sc/ | N/A |
| Biorender | https://www.biorender.com/ | N/A |

## EXPERIMENTAL MODEL AND STUDY PARTICIPANT DETAILS

### Cell lines

Human THP-1 monocytes were cultured in T75 flasks in RPMI 1640 medium supplemented with 10% fetal bovine serum (FBS), 2 mM L-glutamine, and 100 U/mL penicillin-streptomycin, in a humidified incubator at 37 °C with 5% CO₂. For differentiation into macrophage-like cells, cells were treated with 200 ng/mL phorbol 12-myristate 13-acetate (PMA; Sigma-Aldrich) for 3 hours at 37 °C. After treatment, PMA was removed by centrifugation at 280 × g for 5 minutes, and cells were resuspended in fresh medium and seeded into 96-well plates at a density of 1 × 10⁵ cells per well one day prior to experiments.^50^ These differentiated cells were used in all subsequent functional experiments. Murine alveolar macrophage (AM)-like cells were generated from fetal liver cells in the presence of GM-CSF as previously described.^53^ Cells were cultured in T75 flasks in RPMI 1640 supplemented with 10% FBS, 2 mM L-glutamine, and 10 mM HEPES at 37 °C and 5% CO₂. The culture medium was supplemented with 30 ng/mL GM-CSF (BioLegend). Upon reaching ∼90% confluence, cells were detached with 1.5 mM EDTA in DPBS for 3 min at 37 °C, collected by centrifugation (280 × g, 3 min), resuspended in fresh medium, and seeded into 96-well plates at 1 × 10⁵ cells per well.

### Mice

Wild-type and *Nlrp3*^-/-^ mice ^102^ on a C57BL/6 background were housed under SPF conditions at the Center for Experimental Models and Transgenic Services (CEMT), Freiburg, Germany, in accordance with local guidelines.

### BMDM and BMDC preparation

Murine bone marrow-derived macrophages (BMDMs) and bone marrow-derived dendritic cells (BMDCs) were differentiated from tibial and femoral bone marrow as previously described in detail.^103^ Cells were cultured at 37 °C with 5% CO₂ in a humidified incubator in RPMI 1640 medium containing GlutaMAX (Life Technologies GmbH), supplemented with 10% FCS, 100 U/mL penicillin-streptomycin, and either 20 ng/mL recombinant murine GM-CSF (Immunotools) for BMDC differentiation or 100 ng/mL recombinant human M-CSF (7BioScience GmbH) for BMDM differentiation. After 6–8 days of differentiation, cells were plated in 96-well plates at a density of 0.07–0.08 × 10^6^ cells per well the day before the experiments.

### PBMC-derived monocytes preparation

Human peripheral blood mononuclear cells (PBMCs) were isolated from venous blood of healthy male and female donors with a median age of 38 years by density gradient centrifugation with a Ficoll-Paque PLUS gradient (VWR International GmbH). Isolated cells were cultured at 37 °C and 5% CO_2_. The cells were rested for 2 hours after isolation in RPMI containing GlutaMAX without supplements. Non-adherent cells were washed away and cells were rested in BMDM medium overnight before being used for experiments. Then, non-adherent cells were washed away and fresh BMDM medium was added. The adherent cells were used as PBMC-derived monocytes for subsequent experiments.

### Bacterial strains and culture

*P. aeruginosa* PAO1 and *E. coli* BL21(DE3) transformed with the plasmid pET25PAIIL (based on the pET25b (+) vector), were cultivated overnight for 16 hours in Lisogeny Broth (LB) medium at 37 °C and shaking at 180 rpm. The *E. coli* was grown in the presence of 100 μg/ml ampicillin. The grown culture of transformed *E. coli* was diluted down to OD_600_ = 0.8 and with LB medium containing the antibiotic and 1mM isopropyl β-D-1-thiogalactopyranoside (IPTG; Carl Roth) and incubated at 30 °C with the shaking at 180 rpm to induce LecB expression. Before infection, bacterial cultures were diluted in PBS to the concentration required to achieve the indicated multiplicity of infection (MOI), based on colony-forming units (CFU). Where indicated, bacteria were preincubated with the respective sugar inhibitor for 30 minutes at room temperature with gentle shaking.

## METHOD DETAILS

### Production of LecA and LecB

Recombinant *P. aeruginosa* LecA and LecB were expressed in *E. coli* BL21(DE3) cells transformed with plasmids pET25PAIL and pET25PAIIL, respectively (both based on the pET25b(+) vector), and purified by affinity chromatography. LecA was purified using D-galactose agarose (Thermo Fisher Scientific), and LecB was purified using D-mannose agarose (Sigma-Aldrich). The purity of the recombinant proteins was >98%, as assessed by SDS-PAGE. Proteins were extensively dialyzed for 7 days to remove residual sugars. ^104,105^ Final preparations were flash-frozen in liquid nitrogen and stored at −80 °C, as previously described.

### Cell stimulation

THP-1 cells and AM-like cells were primed with 150 ng/mL lipopolysaccharide (LPS; Sigma-Aldrich) for 3 hours. BMDCs, BMDMs, and PBMC-derived monocytes were primed with 20 ng/mL LPS (InvivoGen) for 3 hours. Where indicated, cells were pretreated with specific inhibitors for 30 minutes prior to stimulation, and inhibitors were maintained throughout the subsequent stimulation period. THP-1 cells and AM-like cells were treated with MCC950 (3 μM; Adipogen), KCl (50 mM; Carl Roth), or Ebselen (40 μM; Sigma-Aldrich), whereas BMDMs, BMDCs, and PBMC-derived monocytes were treated with MCC950 (5 μM; Adipogen), KCl (60 mM; Sigma-Aldrich), or Ebselen (30 μM; Biomol GmbH). THP-1 cells were also treated with RGD peptide (100 or 500 μM; MedChemExpress) or an anti-integrin α5β1 antibody (70 nM; Sigma-Aldrich). For glycosphingolipid depletion experiments, AM-like cells were pretreated with PPMP (2.5 μM; Santa Cruz) for 4 days, with daily replacement of PPMP-containing medium. PPMP was maintained throughout the subsequent stimulation period. Depletion of cell-surface globotriaosylceramide was assessed by immunofluorescence in binding assays using Shiga toxin B subunit (StxB; Sigma-Aldrich) conjugated to Alexa Fluor 488 (StxB-AF488; 0.25 μM). Unless otherwise indicated, LecA or LecB was used at 0.5 μM for 24 hours. Nigericin (Sigma-Aldrich) was used at 5 μM for 1–3 hours depending on the cell type and experiment. For carbohydrate competition experiments, LecB was preincubated with L-fucose (45 mM; Sigma-Aldrich) and LecA with PNPG (10 mM; Sigma-Aldrich) for 30 minutes at room temperature before application to cells.

### Western blotting

For immunoblot analysis, cell culture plates were centrifuged at 400 × g for 5 minutes at 4 °C. The supernatant was carefully collected without disturbing the cell layer and mixed with an appropriate volume of 3× sample buffer (187.5 mM Tris-HCl, pH 6.8; 6% SDS; 30% glycerol; 0.03% phenol red) containing 66 mM DTT.^106^ The remaining supernatant was discarded, and the adherent cells were washed once with cold PBS, followed by centrifugation at 400 × g for 5 minutes at 4 °C. After removing the wash buffer, cells were lysed in 1× sample buffer (prepared by diluting 3× sample buffer with ddH_2_O), vortexed, and heated at 98 °C for 8 minutes. Protein samples were separated by SDS-PAGE and transferred to nitrocellulose membranes. Membranes were blocked with 2% milk in PBST (PBS + 0.1% Tween-20) for 1 hour at room temperature or overnight at 4 °C. The membranes were then incubated with primary antibodies for 1–2 hours at room temperature or overnight at 4 °C.^103^ The following primary antibodies were used: Anti-Caspase-1 (p20) (human) mAb (Bally-1, Adipogen; 1:1000), Anti-Caspase-1 (p20) (mouse) mAb (Casper-1, Adipogen; 1:1000), Anti-NLRP3/NALP3 mAb (Cryo-2, Adipogen; 1:1000), Anti-IL-1β (D3U3E, Cell Signaling; 1:800), Mouse IL-1β/IL-1F2 Antibody (R&D Systems; 1:800), anti-GSDMD (126-138, Sigma-Aldrich; 1:1000), Anti-GSDMD antibody (EPR19828, Abcam; 1:500), Mouse anti-integrin β1 (Millipore; 1:1000), Anti-LecB (Eurogentec; 1:500), and Anti-beta-Actin (Sigma-Aldrich; 1:2000). After incubation with primary antibodies, membranes were washed three times with PBST (5 minutes each), incubated with HRP-conjugated secondary antibodies (Cell Signaling; 1:2000) for 1 hour at room temperature, washed again as above, and visualized using either the Novex™ ECL Chemiluminescent Substrate Reagent Kit (Thermo Fisher Scientific) or the Westar Supernova substrate (Cyanagen).

### ELISA

IL-1β levels were quantified in cell-free culture supernatants using ELISA kits (Thermo Fisher Scientific), following the manufacturer’s instructions. Briefly, 96-well ELISA plates were coated with 100 μL of IL-1β capture antibody solution and incubated overnight at 4 °C. The next day, plates were blocked with 1× ELISA/ELISPOT diluent for 1 hour at room temperature. Standards and samples were diluted in 1× ELISA/ELISPOT diluent and added to the wells, followed by incubation for 2 hours at room temperature or overnight at 4 °C. After washing, detection antibody was added and incubated for 1 hour at room temperature, followed by incubation with streptavidin-HRP for 30 minutes. Plates were developed using 1× TMB substrate solution for 15 minutes, and the reaction was stopped by adding 2 N sulfuric acid. Absorbance was measured at 450 nm using a microplate reader.

### LDH release assay

Lactate dehydrogenase (LDH) release in cell-free supernatants was quantified using the CytoTox 96® Non-Radioactive Cytotoxicity Assay (Promega) according to the manufacturer’s instructions. Cell-free culture medium served as a blank control, and values were background-subtracted accordingly. Maximum LDH release (100% lysis) was determined by treating cells with lysis buffer 45 minutes before supernatant collection. After addition of the CytoTox 96® Reagent, plates were incubated at room temperature for 30 minutes. The reaction was stopped with stop solution, and absorbance was measured at 490 nm within 1 hour using a microplate reader. Data are expressed as the percentage of maximum LDH release.

### Fluorescent labeling of lectins

LecA and LecB were labeled with Alexa Fluor™ 488 NHS-Ester (Succinimidylester) (Thermo Fisher Scientific). Briefly, 10 μL of 1 M NaHCO₃ was added per 100 μL of protein solution, followed by incubation with the fluorophore (dissolved in DMSO) at room temperature for 1 hour with continuous stirring. Unreacted dye was removed using Zeba™ Spin Desalting Columns (7 kDa MWCO; Thermo Fisher Scientific) by centrifugation at 1,500 × g for 2 minutes. The labeled proteins were collected and stored in the dark at 4 °C. Protein concentration was determined using a NanoDrop 2000 spectrophotometer (Thermo Fisher Scientific).

### Biotinylation of LecB

LecB was biotinylated using the EZ-Link NHS-PEG4 Biotinylation Kit (Thermo Fisher Scientific). EZ-Link NHS-PEG4-Biotin was freshly dissolved in 170 µL ddH_2_O and immediately added to the lectin solution at a 20-fold molar excess, followed by incubation for 2 hours at 4 °C. Unreacted biotinylation reagent was removed using Zeba Spin Desalting Columns (7 kDa MWCO) by centrifugation at 1,000 × g for 2 minutes. The degree of biotinylation of LecB was determined using a HABA/avidin assay (A_500_).

### Pull-down assay

THP-1 cells were incubated with LecB-biotin as indicated and lysed in lysis buffer (100 mM Tris-HCl, pH 8.0; 150 mM NaCl; 1 mM EDTA; 1% IGEPAL; 5% glycerol) supplemented with protease inhibitors. Lysate was clarified by centrifugation (17,000 × g, 10 minutes, 4 °C). Streptavidin magnetic beads (Pierce™, Thermo Fisher Scientific) were pre-washed with binding/wash buffer (TBS containing 0.1% Tween-20) by gentle inversion. For each sample, 50 µL of beads were incubated with 180 µL of clarified lysate for 1 hour at room temperature with mixing. After magnetic separation, an aliquot of the lysate was retained as input and the unbound fraction was collected as flow-through. Beads were washed three times with binding/wash buffer, and bound proteins were eluted by adding 1× SDS sample buffer (prepared by diluting 5× SDS sample buffer with ddH_2_O) and heating at 95 °C for 10 minutes. Input and flow-through fractions were also mixed with 5× SDS sample buffer (250 mM Tris-HCl, pH 6.8; 10% SDS; 30% glycerol; 5% β-mercaptoethanol; 0.02% bromophenol blue) and heated at 95 °C for 10 minutes. Samples were analyzed by SDS-PAGE and western blotting.

### Lectin blotting assay (PNGase F digestion)

THP-1 cells were lysed in LecB lysis buffer (100 mM Tris-HCl, pH 8.0; 150 mM NaCl; 1 mM EDTA; 1% IGEPAL; 5% glycerol) supplemented with protease inhibitors. Lysate was clarified by centrifugation (17,000 × g, 10 minutes, 4 °C), and supernatant was transferred to fresh tubes. Denaturation buffer (0.2% SDS, 100 mM β-mercaptoethanol) was added to the lysate at a 1:10 (v/v) ratio, followed by incubation at 100 °C for 10 minutes. Samples were cooled on ice and treated with PNGase F (37 °C, 3 hours; Sigma-Aldrich); matched samples without PNGase F served as controls. Reactions were stopped by adding 5× SDS sample buffer and heating at 100 °C for 5 minutes. Samples were resolved by SDS-PAGE (8% gel) and transferred to nitrocellulose membranes. Membranes were blocked in protein-free blocking buffer (LI-COR) for 1 hour at room temperature and then incubated overnight at 4 °C with LecB-AF488 (1:70 dilution of a ∼1.5 mg/mL stock) in blocking buffer supplemented with 1 mM CaCl_2_ and 1 mM MgCl_2_. Membranes were washed three times with TBST (supplemented with 1 mM CaCl₂ and 1 mM MgCl₂) and imaged using a fluorescence imager. For immunoblotting, membranes were probed with anti-integrin β1 (Millipore; 1:1000) followed by goat anti-mouse Alexa Fluor 647-conjugated secondary antibody (Thermo Fisher Scientific; 1:1000), washed with TBST, and imaged.

### Flow cytometry

THP-1 cells and AM-like cells were seeded at a density of 1 × 10⁵ cells per well in 96-well U-bottom plates and centrifuged at 1,600 × g for 3 minutes at 4 °C. Cells were resuspended in lectin-containing medium, with or without inhibitors, and incubated on ice for 30 minutes, as specified in the figure legends. After incubation, cells were washed with FACS buffer (PBS supplemented with 3% FBS). Flow cytometry was performed using an Attune NxT flow cytometer (Thermo Fisher Scientific, Waltham, MA, USA), and data were analyzed using FlowJo software (BD Biosciences).

### Confocal microscopy

THP-1 cells and AM-like cells were seeded at a density of 5 × 10⁵ cells per well onto 12 mm glass coverslips placed in 4-well plates and allowed to adhere overnight. Cells were then treated with lectins (including Alexa Fluor 488-labeled LecA or LecB) or inhibitors for the indicated time, as specified in the figure legends. Following stimulation, cells were washed three times with PBS and fixed with 4% paraformaldehyde (PFA) for 15 minutes at room temperature. Fixation was followed by quenching with 50 mM ammonium chloride (NH₄Cl) for 5 – 10 minutes. Cells were then permeabilized and blocked in PBS containing 0.2% saponin and 3% BSA for 30 minutes at room temperature.

For ASC staining, cells were incubated with anti-ASC antibody (Adipogen; 1:100 in 3% BSA) for 1 hour at room temperature, followed by PBS washes and incubation with goat anti-rabbit Alexa Fluor 647-conjugated secondary antibody (Thermo Fisher Scientific; 1:200) and DAPI (Carl Roth; 1:500) for 25 minutes at room temperature.

For NLRP3 staining, cells were incubated with anti-NLRP3 antibody (Adipogen; 1:100 in 3% BSA) for 1 hour, followed by goat anti-mouse Alexa Fluor 647-conjugated secondary antibody (Thermo Fisher Scientific; 1:200) and DAPI (Carl Roth; 1:500) under the same conditions.

Samples were dried and imaged using a Nikon A1R confocal microscope equipped with a 60× oil immersion objective (NA 1.49). Laser wavelengths of 405 nm, 488 nm, and 647 nm were used for fluorescence excitation. Images were acquired and processed using ImageJ Fiji software.

### Cell harvesting and lipid extraction for LC-MS analysis

THP-1 cells and AM-like cells were harvested as previously described.^107^ Briefly, cells were washed twice with 0.9% NaCl and quenched with 1 mL of ice-cold methanol:water (1:1, v/v). Cells were scraped, transferred to a new vial, snap-frozen in liquid nitrogen, and stored at −80 °C until further processing.

Samples were thawed on ice, supplemented with 500 μL chloroform, and shaken for 10 minutes at 1,400 rpm and 4 °C. After centrifugation for 10 minutes at 20,000 × g and 4 °C, 200 μL of the chloroform phase was transferred to a new vial and evaporated. The lipid pellets were resuspended in 100 μL isopropanol:acetonitrile:water (2:1:1, v/v/v). Targeted lipid profiling by LC-QqQ-MS was conducted as described previously.^108^

For identification of unknown lipid features, the same samples were subjected to high-resolution mass spectrometry using a Synapt G2-Si mass spectrometer (Waters). The parameters were set as follows: capillary voltage, 3 kV; source temperature, 140 °C; sampling cone voltage, 80 V; source offset, 40 V; desolvation gas flow, 900 L/h at 500 °C; cone gas flow, 50 L/h; and nebulizer pressure, 6.5 bar. Samples were analyzed in positive ion mode using full-scan MS and MS/MS acquisition.

## QUANTIFICATION AND STATISTICAL ANALYSIS

Statistical details, including the type of test used and the exact sample size (n), are provided in the corresponding figure legends. Data are presented as mean ± standard deviation (SD). The number of biological replicates is indicated in each figure legend. Statistical analyses were performed using GraphPad Prism software. Significance was determined as follows: *P* < 0.05 (*), *P* < 0.01 (**), *P* < 0.001 (***), *P* < 0.0001 (****); ns, not significant.

### Declaration of generative AI and AI-assisted technologies in the manuscript preparation process

During the preparation of this manuscript, the authors used ChatGPT (OpenAI) to assist with language editing, improving clarity, and checking the logical flow of the text. The authors reviewed and edited the output as needed and take full responsibility for the content of the manuscript.

## REFERENCES

1. Azam, M.W., and Khan, A.U. (2019). Updates on the pathogenicity status of Pseudomonas aeruginosa. Drug Discovery Today 24, 350–359. 10.1016/j.drudis.2018.07.003.

2. Tarazi, Y.H., Abu-Basha, E., Ismail, Z.B., and Al-Jawasreh, S.I. (2021). Antimicrobial susceptibility of multidrug-resistant Pseudomonas aeruginosa isolated from drinking water and hospitalized patients in Jordan. Acta Tropica 217, 105859. 10.1016/j.actatropica.2021.105859.

3. Arai, H. (2011). Regulation and Function of Versatile Aerobic and Anaerobic Respiratory Metabolism in Pseudomonas aeruginosa. Front. Microbio. 2. 10.3389/fmicb.2011.00103.

4. Qin, S., Xiao, W., Zhou, C., Pu, Q., Deng, X., Lan, L., Liang, H., Song, X., and Wu, M. (2022). Pseudomonas aeruginosa: pathogenesis, virulence factors, antibiotic resistance, interaction with host, technology advances and emerging therapeutics. Sig Transduct Target Ther 7. 10.1038/s41392-022-01056-1.

5. Liao, C., Huang, X., Wang, Q., Yao, D., and Lu, W. (2022). Virulence Factors of Pseudomonas Aeruginosa and Antivirulence Strategies to Combat Its Drug Resistance. Front. Cell. Infect. Microbiol. 12. 10.3389/fcimb.2022.926758.

6. Sati, H., Carrara, E., Savoldi, A., Hansen, P., Garlasco, J., Campagnaro, E., Boccia, S., Castillo-Polo, J.A., Magrini, E., Garcia-Vello, P., et al. (2025). The WHO Bacterial Priority Pathogens List 2024: a prioritisation study to guide research, development, and public health strategies against antimicrobial resistance. The Lancet Infectious Diseases. 10.1016/s1473-3099(25)00118-5.

7. Barken, K.B., Pamp, S.J., Yang, L., Gjermansen, M., Bertrand, J.J., Klausen, M., Givskov, M., Whitchurch, C.B., Engel, J.N., and Tolker-Nielsen, T. (2008). Roles of type IV pili, flagellum-mediated motility and extracellular DNA in the formation of mature multicellular structures in *Pseudomonas aeruginosa* biofilms. Environmental Microbiology 10, 2331–2343. 10.1111/j.1462-2920.2008.01658.x.

8. Hahn, H.P. (1997). The type-4 pilus is the major virulence-associated adhesin of Pseudomonas aeruginosa – a review. Gene 192, 99–108. 10.1016/s0378-1119(97)00116-9.

9. Garcia, M., Morello, E., Garnier, J., Barrault, C., Garnier, M., Burucoa, C., Lecron, J.-C., Si-Tahar, M., Bernard, F.-X., and Bodet, C. (2018). *Pseudomonas aeruginosa* flagellum is critical for invasion, cutaneous persistence and induction of inflammatory response of skin epidermis. Virulence 9, 1163–1175. 10.1080/21505594.2018.1480830.

10. Jouault, A., Saliba, A.M., and Toueui, L. (2022). Modulation of the immune response by the Pseudomonas aeruginosa type-III secretion system. Front. Cell. Infect. Microbiol. 12. 10.3389/fcimb.2022.1064010.

11. Hauser, A.R. (2009). The type III secretion system of Pseudomonas aeruginosa: infection by injection. Nat Rev Microbiol 7, 654–665. 10.1038/nrmicro2199.

12. Horna, G., and Ruiz, J. (2021). Type 3 secretion system of Pseudomonas aeruginosa. Microbiological Research 246, 126719. 10.1016/j.micres.2021.126719.

13. Gheorghita, A.A., Wozniak, D.J., Parsek, M.R., and Howell, P.L. (2023). *Pseudomonas aeruginosa* biofilm exopolysaccharides: assembly, function, and degradation. FEMS Microbiology Reviews 47. 10.1093/femsre/fuad060.

14. Li, Q., Mao, S., Wang, H., and Ye, X. (2022). The Molecular Architecture of Pseudomonas aeruginosa Quorum-Sensing Inhibitors. Marine Drugs 20, 488. 10.3390/md20080488.

15. Singh, V.K., Almpani, M., Wheeler, K.M., and Rahme, L.G. (2023). Interconnections of Pseudomonas aeruginosa Quorum-Sensing Systems in Intestinal Permeability and Inflammation. mBio 14. 10.1128/mbio.03524-22.

16. Imberty, A., Wimmerová, M., Mitchell, E.P., and Gilboa-Garber, N. (2004). Structures of the lectins from Pseudomonas aeruginosa: insights into the molecular basis for host glycan recognition. Microbes and Infection 6, 221–228. 10.1016/j.micinf.2003.10.016.

17. Grishin, A.V., Krivozubov, M.S., Karyagina, A.S., and Gintsburg, A.L. (2015). Pseudomonas Aeruginosa Lectins As Targets for Novel Antibacterials. Acta Naturae 7, 29–41. 10.32607/20758251-2015-7-2-29-41.

18. Faltinek, L., Melicher, F., Kelemen, V., Mező, E., Borbás, A., and Wimmerová, M. (2025). Bispecific Thio-Linked Disaccharides as Inhibitors of *Pseudomonas Aeruginosa* Lectins LecA (PA-IL) and LecB (PA-IIL): Dual-Targeting Strategy. Chemistry A European J 31. 10.1002/chem.202403546.

19. Siukstaite, L., Imberty, A., and Römer, W. (2021). Structural Diversities of Lectins Binding to the Glycosphingolipid Gb3. Front. Mol. Biosci. 8, 704685. 10.3389/fmolb.2021.704685.

20. Omidvar, R., Ayala, Y.A., Brandel, A., Hasenclever, L., Helmstädter, M., Rohrbach, A., Römer, W., and Madl, J. (2021). Quantification of nanoscale forces in lectin-mediated bacterial attachment and uptake into giant liposomes. Nanoscale 13, 4016–4028. 10.1039/d0nr07726g.

21. Eierhoff, T., Bastian, B., Thuenauer, R., Madl, J., Audfray, A., Aigal, S., Juillot, S., Rydell, G.E., Müller, S., De Bentzmann, S., et al. (2014). A lipid zipper triggers bacterial invasion. Proc. Natl. Acad. Sci. U.S.A. 111, 12895–12900. 10.1073/pnas.1402637111.

22. Cioci, G., Mitchell, E.P., Gautier, C., Wimmerová, M., Sudakevitz, D., Pérez, S., Gilboa-Garber, N., and Imberty, A. (2003). Structural basis of calcium and galactose recognition by the lectin PA-IL of *Pseudomonas aeruginosa*. FEBS Letters 555, 297–301. 10.1016/s0014-5793(03)01249-3.

23. Zheng, S., Eierhoff, T., Aigal, S., Brandel, A., Thuenauer, R., De Bentzmann, S., Imberty, A., and Römer, W. (2017). The Pseudomonas aeruginosa lectin LecA triggers host cell signalling by glycosphingolipid-dependent phosphorylation of the adaptor protein CrkII. Biochimica et Biophysica Acta (BBA) - Molecular Cell Research 1864, 1236–1245. 10.1016/j.bbamcr.2017.04.005.

24. Brandel, A., Aigal, S., Lagies, S., Schlimpert, M., Meléndez, A.V., Xu, M., Lehmann, A., Hummel, D., Fisch, D., Madl, J., et al. (2021). The Gb3-enriched CD59/flotillin plasma membrane domain regulates host cell invasion by Pseudomonas aeruginosa. Cell. Mol. Life Sci. 78, 3637–3656. 10.1007/s00018-021-03766-1.

25. Aigal, S., Omidvar, R., Stober, K., Ziegelbauer, J., Eierhoff, T., Schampera, J.N., Römer, W., and Schwan, C. (2022). Septin barriers protect mammalian host cells against Pseudomonas aeruginosa invasion. Cell Reports 41, 111510. 10.1016/j.celrep.2022.111510.

26. Avichezer, D., and Gilboa-Garber, N. (1987). PA-II, the L-fucose and D-mannose binding lectin of *Pseudomonas aeruginosa* stimulates human peripheral lymphocytes and murine splenocytes. FEBS Letters 216, 62–66. 10.1016/0014-5793(87)80757-3.

27. Garber, N., Guempel, U., Gilboa-Garber, N., and Royle, R.J. (1987). Specificity of the fucose-binding lectin of*Pseudomonas aeruginosa*. FEMS Microbiology Letters 48, 331– 334. 10.1111/j.1574-6968.1987.tb02619.x.

28. Mitchell, E.P., Sabin, C., Šnajdrová, L., Pokorná, M., Perret, S., Gautier, C., Hofr, C., Gilboa-Garber, N., Koča, J., Wimmerová, M., et al. (2005). High affinity fucose binding of *Pseudomonas aeruginosa* lectin PA-IIL: 1.0 Å resolution crystal structure of the complex combined with thermodynamics and computational chemistry approaches. Proteins 58, 735–746. 10.1002/prot.20330.

29. Sabin, C., Mitchell, E.P., Pokorná, M., Gautier, C., Utille, J.-P., Wimmerová, M., and Imberty, A. (2006). Binding of different monosaccharides by lectin PA-IIL from *Pseudomonas aeruginosa*: Thermodynamics data correlated with X-ray structures. FEBS Letters 580, 982–987. 10.1016/j.febslet.2006.01.030.

30. Hofmann, A., Sommer, R., Hauck, D., Stifel, J., Göttker-Schnetmann, I., and Titz, A. (2015). Synthesis of mannoheptose derivatives and their evaluation as inhibitors of the lectin LecB from the opportunistic pathogen Pseudomonas aeruginosa. Carbohydrate Research 412, 34–42. 10.1016/j.carres.2015.04.010.

31. Passos Da Silva, D., Matwichuk, M.L., Townsend, D.O., Reichhardt, C., Lamba, D., Wozniak, D.J., and Parsek, M.R. (2019). The Pseudomonas aeruginosa lectin LecB binds to the exopolysaccharide Psl and stabilizes the biofilm matrix. Nat Commun 10. 10.1038/s41467-019-10201-4.

32. Cott, C., Thuenauer, R., Landi, A., Kühn, K., Juillot, S., Imberty, A., Madl, J., Eierhoff, T., and Römer, W. (2016). Pseudomonas aeruginosa lectin LecB inhibits tissue repair processes by triggering β-catenin degradation. Biochimica et Biophysica Acta (BBA) - Molecular Cell Research 1863, 1106–1118. 10.1016/j.bbamcr.2016.02.004.

33. Landi, A., Mari, M., Kleiser, S., Wolf, T., Gretzmeier, C., Wilhelm, I., Kiritsi, D., Thünauer, R., Geiger, R., Nyström, A., et al. (2019). *Pseudomonas aeruginosa*lectin LecB impairs keratinocyte fitness by abrogating growth factor signalling. Life Sci. Alliance 2, e201900422. 10.26508/lsa.201900422.

34. Thuenauer, R., Landi, A., Trefzer, A., Altmann, S., Wehrum, S., Eierhoff, T., Diedrich, B., Dengjel, J., Nyström, A., Imberty, A., et al. (2020). The Pseudomonas aeruginosa Lectin LecB Causes Integrin Internalization and Inhibits Epithelial Wound Healing. mBio 11. 10.1128/mbio.03260-19.

35. Thuenauer, R., Kühn, K., Guo, Y., Kotsis, F., Xu, M., Trefzer, A., Altmann, S., Wehrum, S., Heshmatpour, N., Faust, B., et al. (2022). The Lectin LecB Induces Patches with Basolateral Characteristics at the Apical Membrane to Promote Pseudomonas aeruginosa Host Cell Invasion. mBio 13, e00819–22. 10.1128/mbio.00819-22.

36. Sponsel, J., Guo, Y., Hamzam, L., Lavanant, A.C., Pérez-Riverón, A., Partiot, E., Muller, Q., Rottura, J., Gaudin, R., Hauck, D., et al. (2023). *Pseudomonas aeruginosa* LECB suppresses immune responses by inhibiting transendothelial migration. EMBO Reports 24, e55971. 10.15252/embr.202255971.

37. Amarante-Mendes, G.P., Adjemian, S., Branco, L.M., Zanetti, L.C., Weinlich, R., and Bortoluci, K.R. (2018). Pattern Recognition Receptors and the Host Cell Death Molecular Machinery. Front. Immunol. 9. 10.3389/fimmu.2018.02379.

38. Chen, R., Zou, J., Chen, J., Zhong, X., Kang, R., and Tang, D. (2025). Pattern recognition receptors: function, regulation and therapeutic potential. Sig Transduct Target Ther 10. 10.1038/s41392-025-02264-1.

39. Akira, S., Uematsu, S., and Takeuchi, O. (2006). Pathogen Recognition and Innate Immunity. Cell 124, 783–801. 10.1016/j.cell.2006.02.015.

40. Medzhitov, R. (2007). Recognition of microorganisms and activation of the immune response. Nature 449, 819–826. 10.1038/nature06246.

41. Schroder, K., and Tschopp, J. (2010). The Inflammasomes. Cell 140, 821–832. 10.1016/j.cell.2010.01.040.

42. Li, D., and Wu, M. (2021). Pattern recognition receptors in health and diseases. Sig Transduct Target Ther 6, 291. 10.1038/s41392-021-00687-0.

43. Broz, P., and Dixit, V.M. (2016). Inflammasomes: mechanism of assembly, regulation and signalling. Nat Rev Immunol 16, 407–420. 10.1038/nri.2016.58.

44. Zheng, D., Liwinski, T., and Elinav, E. (2020). Inflammasome activation and regulation: toward a better understanding of complex mechanisms. Cell Discov 6, 36. 10.1038/s41421-020-0167-x.

45. Sharma, M., and De Alba, E. (2021). Structure, Activation and Regulation of NLRP3 and AIM2 Inflammasomes. IJMS 22, 872. 10.3390/ijms22020872.

46. Liu, F., Shao, B., Zhu, Y., Xue, X., and Wu, X. (2025). Mechanisms of NLRP3 inflammasome in pathogenesis and progression of inflammation-related gastrointestinal diseases. PeerJ 13, e19828. 10.7717/peerj.19828.

47. Thompson, M., Williams, J., Anderson, D., and Martin, E. (2025). NLRP3 inflammasome activation pathways and their contribution to the pathophysiology of chronic inflammatory and metabolic diseases. Int. J. Adv. Biochem. Res. 9, 73–78. 10.33545/26174693.2025.v9.i7a.7564.

48. Minns, M.S., Liboro, K., Lima, T.S., Abbondante, S., Miller, B.A., Marshall, M.E., Tran Chau, J., Roistacher, A., Rietsch, A., Dubyak, G.R., et al. (2023). NLRP3 selectively drives IL-1β secretion by Pseudomonas aeruginosa infected neutrophils and regulates corneal disease severity. Nat Commun 14, 5832. 10.1038/s41467-023-41391-7.

49. Ge, J., Cao, M., Zhang, Y., Wu, T., Liu, J., Pu, J., He, H., Guo, Z., Ju, S., and Yu, J. (2025). Inhibiting NLRP3 enhances cellular autophagy induced by outer membrane vesicles from *Pseudomonas aeruginosa*. Microbiol Spectr 13. 10.1128/spectrum.01819-24.

50. Saller, B.S., Neuwirt, E., and Groß, O. (2023). Methods to Activate the NLRP3 Inflammasome. In NLR Proteins Methods in Molecular Biology., P. Pelegrjn and F. Di Virgilio, eds. (Springer US), pp. 169–197. 10.1007/978-1-0716-3350-2_12.

51. Chanput, W., Mes, J.J., and Wichers, H.J. (2014). THP-1 cell line: An in vitro cell model for immune modulation approach. International Immunopharmacology 23, 37–45. 10.1016/j.intimp.2014.08.002.

52. Shah, P.T., Tufail, M., Wu, C., and Xing, L. (2022). THP-1 cell line model for tuberculosis: A platform for in vitro macrophage manipulation. Tuberculosis 136, 102243. 10.1016/j.tube.2022.102243.

53. Fejer, G., Wegner, M.D., Györy, I., Cohen, I., Engelhard, P., Voronov, E., Manke, T., Ruzsics, Z., Dölken, L., Prazeres Da Costa, O., et al. (2013). Nontransformed, GM-CSF– dependent macrophage lines are a unieue model to study tissue macrophage functions. Proc. Natl. Acad. Sci. U.S.A. 110. 10.1073/pnas.1302877110.

54. Blanchard, B., Nurisso, A., Hollville, E., Tétaud, C., Wiels, J., Pokorná, M., Wimmerová, M., Varrot, A., and Imberty, A. (2008). Structural Basis of the Preferential Binding for Globo-Series Glycosphingolipids Displayed by Pseudomonas aeruginosa Lectin I. Journal of Molecular Biology 383, 837–853. 10.1016/j.jmb.2008.08.028.

55. Abe, A., Inokuchi, J., Jimbo, M., Shimeno, H., Nagamatsu, A., Shayman, J.A., Shukla, G.S., and Radin, N.S. (1992). Improved Inhibitors of Glucosylceramide Synthase1. The Journal of Biochemistry 111, 191–196. 10.1093/oxfordjournals.jbchem.a123736.

56. Mahfoud, R., Manis, A., and Lingwood, C.A. (2009). Fatty acid-dependent globotriaosyl ceramide receptor function in detergent resistant model membranes. Journal of Lipid Research 50, 1744–1755. 10.1194/jlr.M800385-JLR200.

57. Deng, Q., Wang, Y., Zhang, Y., Li, M., Li, D., Huang, X., Wu, Y., Pu, J., and Wu, M. (2016). Pseudomonas aeruginosa Triggers Macrophage Autophagy To Escape Intracellular Killing by Activation of the NLRP3 Inflammasome. Infect Immun 84, 56–66. 10.1128/iai.00945-15.

58. Tan, Q., Ai, Q., He, Y., Li, F., and Yu, J. (2022). P. aeruginosa biofilm activates the NLRP3 inflammasomes in vitro. Microbial Pathogenesis 164, 105379. 10.1016/j.micpath.2021.105379.

59. White, A., Wang, Z., Wang, X., King, M., Guo, C., Mantsounga, C., Ayala, A., Morrison, A.R., Choudhary, G., Sellke, F., et al. (2022). NLRP3 inflammasome activation in cigarette smoke priming for Pseudomonas aeruginosa-induced acute lung injury. Redox Biology 57, 102467. 10.1016/j.redox.2022.102467.

60. Muñoz-Planillo, R., Kuffa, P., Martjnez-Colón, G., Smith, B.L., Rajendiran, T.M., and Núñez, G. (2013). K+ Efflux Is the Common Trigger of NLRP3 Inflammasome Activation by Bacterial Toxins and Particulate Matter. Immunity 38, 1142–1153. 10.1016/j.immuni.2013.05.016.

61. Zhou, R., Yazdi, A.S., Menu, P., and Tschopp, J. (2011). A role for mitochondria in NLRP3 inflammasome activation. Nature 469, 221–225. 10.1038/nature09663.

62. Groß, C.J., Mishra, R., Schneider, K.S., Médard, G., Wettmarshausen, J., Dittlein, D.C., Shi, H., Gorka, O., Koenig, P.-A., Fromm, S., et al. (2016). K + Efflux-Independent NLRP3 Inflammasome Activation by Small Molecules Targeting Mitochondria. Immunity 45, 761– 773. 10.1016/j.immuni.2016.08.010.

63. Van Opdenbosch, N., Gurung, P., Vande Walle, L., Fossoul, A., Kanneganti, T.-D., and Lamkanfi, M. (2014). Activation of the NLRP1b inflammasome independently of ASC-mediated caspase-1 autoproteolysis and speck formation. Nat Commun 5, 3209. 10.1038/ncomms4209.

64. Broz, P., Von Moltke, J., Jones, J.W., Vance, R.E., and Monack, D.M. (2010). Differential Reeuirement for Caspase-1 Autoproteolysis in Pathogen-Induced Cell Death and Cytokine Processing. Cell Host & Microbe 8, 471–483. 10.1016/j.chom.2010.11.007.

65. Vanaja, S.K., Rathinam, V.A.K., and Fitzgerald, K.A. (2015). Mechanisms of inflammasome activation: recent advances and novel insights. Trends in Cell Biology 25, 308–315. 10.1016/j.tcb.2014.12.009.

66. Lu, A., Magupalli, V.G., Ruan, J., Yin, Q., Atianand, M.K., Vos, M.R., Schröder, G.F., Fitzgerald, K.A., Wu, H., and Egelman, E.H. (2014). Unified Polymerization Mechanism for the Assembly of ASC-Dependent Inflammasomes. Cell 156, 1193–1206. 10.1016/j.cell.2014.02.008.

67. Stutz, A., Kolbe, C.-C., Stahl, R., Horvath, G.L., Franklin, B.S., Van Ray, O., Brinkschulte, R., Geyer, M., Meissner, F., and Latz, E. (2017). NLRP3 inflammasome assembly is regulated by phosphorylation of the pyrin domain. Journal of Experimental Medicine 214, 1725– 1736. 10.1084/jem.20160933.

68. Magupalli, V.G., Negro, R., Tian, Y., Hauenstein, A.V., Di Caprio, G., Skillern, W., Deng, Q., Orning, P., Alam, H.B., Maliga, Z., et al. (2020). HDAC6 mediates an aggresome-like mechanism for NLRP3 and pyrin inflammasome activation. Science 369, eaas8995. 10.1126/science.aas8995.

69. Sun, J., LaRock, D.L., Skowronski, E.A., Kimmey, J.M., Olson, J., Jiang, Z., O’Donoghue, A.J., Nizet, V., and LaRock, C.N. (2020). The Pseudomonas aeruginosa protease LasB directly activates IL-1β. EBioMedicine 60, 102984. 10.1016/j.ebiom.2020.102984.

70. Elfadadny, A., Ragab, R.F., AlHarbi, M., Badshah, F., Ibáñez-Arancibia, E., Farag, A., Hendawy, A.O., De Los Rjos-Escalante, P.R., Aboubakr, M., Zakai, S.A., et al. (2024). Antimicrobial resistance of Pseudomonas aeruginosa: navigating clinical impacts, current resistance trends, and innovations in breaking therapies. Front. Microbiol. 15. 10.3389/fmicb.2024.1374466.

71. Mini, M., and Kumar, P. (2025). Computational and Experimental Validation of 4-Amino-3-Hydroxynaphthalene-1-Sulfonic Acid as a Novel Antibiofilm Agent. European Journal of Pharmacology, 177892. 10.1016/j.ejphar.2025.177892.

72. Da Cruz Nizer, W.S., Inkovskiy, V., Versey, Z., Strempel, N., Cassol, E., and Overhage, J. (2021). Oxidative Stress Response in Pseudomonas aeruginosa. Pathogens 10, 1187. 10.3390/pathogens10091187.

73. Teddy Grandjean, Anne Boucher, Marion Thepaut, Laura Monlezun, Benoit Guery, Eric Faudry, Eric Kipnis, and Rodrigue Dessein (2017). The human NAIP-NLRC4-inflammasome senses the Pseudomonas aeruginosa T3SS inner-rod protein. International Immunology 29, 377–384. 10.1093/intimm/dxx047.

74. Bauer, R., and Rauch, I. (2020). The NAIP/NLRC4 inflammasome in infection and pathology. Molecular Aspects of Medicine 76, 100863. 10.1016/j.mam.2020.100863.

75. Hardy, K.S., Tuckey, A.N., Housley, N.A., Andrews, J., Patel, M., Al-Mehdi, A.-B., Barrington, R.A., Cassel, S.L., Sutterwala, F.S., and Audia, J.P. (2022). The Pseudomonas aeruginosa Type III Secretion System Exoenzyme Effector ExoU Induces Mitochondrial Damage in a Murine Bone Marrow-Derived Macrophage Infection Model. Infect Immun 90, e00470-21. 10.1128/iai.00470-21.

76. Reuven, A.D., Mwaura, B.W., and Bliska, J.B. (2024). ExoS Effector in *Pseudomonas aeruginosa* Hyperactive Type III Secretion System Mutant Promotes Enhanced Plasma Membrane Rupture in Neutrophils. Preprint at Microbiology, 10.1101/2024.01.24.577040 10.1101/2024.01.24.577040.

77. Martjn-González, L., Ramos, B., Huang, Y., Thurston, T., Towers, G., and Lobato-Máreuez, D. (2026). Cell-autonomous immunity as an integrated immune network against bacterial and viral pathogens. Nat Microbiol 11, 2098–2111. 10.1038/s41564-026-02428-x.

78. Dolowschiak, T., Chassin, C., Ben Mkaddem, S., Fuchs, T.M., Weiss, S., Vandewalle, A., and Hornef, M.W. (2010). Potentiation of Epithelial Innate Host Responses by Intercellular Communication. PLoS Pathog 6, e1001194. 10.1371/journal.ppat.1001194.

79. Jastrab, J.B., and Kagan, J.C. (2024). Strategies of bacterial detection by inflammasomes. Cell Chemical Biology 31, 835–850. 10.1016/j.chembiol.2024.03.009.

80. Villringer, S., Madl, J., Sych, T., Manner, C., Imberty, A., and Römer, W. (2018). Lectin-mediated protocell crosslinking to mimic cell-cell junctions and adhesion. Sci Rep 8. 10.1038/s41598-018-20230-6.

81. Tielker, D., Hacker, S., Loris, R., Strathmann, M., Wingender, J., Wilhelm, S., Rosenau, F., and Jaeger, K.-E. (2005). Pseudomonas aeruginosa lectin LecB is located in the outer membrane and is involved in biofilm formation. Microbiology 151, 1313–1323. 10.1099/mic.0.27701-0.

82. Palmioli, A., Sperandeo, P., Polissi, A., and Airoldi, C. (2019). Targeting Bacterial Biofilm: A New LecA Multivalent Ligand with Inhibitory Activity. ChemBioChem 20, 2911–2915. 10.1002/cbic.201900383.

83. Izzati, F.N., Choksi, H., Giuliana, P., Abd-Rabbo, D., Elsaesser, H., Blundell, A., Affe, V., Kannen, V., Jame-Chenarboo, Z., Schmidt, E.N., et al. (2025). Presentation of immunoregulatory sialoglycans on T cells is divergent between mice and humans. Cell Reports 44, 115933. 10.1016/j.celrep.2025.115933.

84. Szczykutowicz, J. (2023). Ligand Recognition by the Macrophage Galactose-Type C-Type Lectin: Self or Non-Self?—A Way to Trick the Host’s Immune System. IJMS 24, 17078. 10.3390/ijms242317078.

85. Toscano, M.A., Bianco, G.A., Ilarregui, J.M., Croci, D.O., Correale, J., Hernandez, J.D., Zwirner, N.W., Poirier, F., Riley, E.M., Baum, L.G., et al. (2007). Differential glycosylation of TH1, TH2 and TH-17 effector cells selectively regulates susceptibility to cell death. Nat Immunol 8, 825–834. 10.1038/ni1482.

86. Mariathasan, S., Weiss, D.S., Newton, K., McBride, J., O’Rourke, K., Roose-Girma, M., Lee, W.P., Weinrauch, Y., Monack, D.M., and Dixit, V.M. (2006). Cryopyrin activates the inflammasome in response to toxins and ATP. Nature 440, 228–232. 10.1038/nature04515.

87. Pétrilli, V., Papin, S., Dostert, C., Mayor, A., Martinon, F., and Tschopp, J. (2007). Activation of the NALP3 inflammasome is triggered by low intracellular potassium concentration. Cell Death Differ 14, 1583–1589. 10.1038/sj.cdd.4402195.

88. Zanoni, I., Tan, Y., Di Gioia, M., Broggi, A., Ruan, J., Shi, J., Donado, C.A., Shao, F., Wu, H., Springstead, J.R., et al. (2016). An endogenous caspase-11 ligand elicits interleukin-1 release from living dendritic cells. Science 352, 1232–1236. 10.1126/science.aaf3036.

89. Lee, M.-S., Kwon, H., Lee, E.-Y., Kim, D.-J., Park, J.-H., Tesh, V.L., Oh, T.-K., and Kim, M.H. (2016). Shiga Toxins Activate the NLRP3 Inflammasome Pathway To Promote Both Production of the Proinflammatory Cytokine Interleukin-1β and Apoptotic Cell Death. Infect Immun 84, 172–186. 10.1128/IAI.01095-15.

90. Orimo, T., Sasaki, I., Hemmi, H., Ozasa, T., Fukuda-Ohta, Y., Ohta, T., Morinaka, M., Kitauchi, M., Yamaguchi, T., Sato, Y., et al. (2019). Cholera toxin B induces interleukin-1β production from resident peritoneal macrophages through the pyrin inflammasome as well as the NLRP3 inflammasome. International Immunology 31, 657–668. 10.1093/intimm/dxz004.

91. Jun, H.-K., Lee, S.-H., Lee, H.-R., and Choi, B.-K. (2012). Integrin α5β1 Activates the NLRP3 Inflammasome by Direct Interaction with a Bacterial Surface Protein. Immunity 36, 755–768. 10.1016/j.immuni.2012.05.002.

92. Kittel, A.-S., Makshakova, O., Hauerwas, M., Edel, N., Knickmeier, N., Tomisch, J., Aljohmani, A., Yildiz, D., Peyronnet, R., and Römer, W. (2025). LecB from Pseudomonas aeruginosa modulates Piezo1 currents and localization in a time-dependent manner. Cell. Mol. Life Sci. 82, 399. 10.1007/s00018-025-05934-z.

93. Shi, Y., Yang, B., Linghu, Y., Liu, Y., Li, H., Xing, Y., Wu, B., Gao, Y., Liao, L., Zheng, Y., et al. (2025). Piezo1 senses hyphae and initiates host defense against pathogenic fungi. Cell Reports 44, 115839. 10.1016/j.celrep.2025.115839.

94. Sych, T., Omidvar, R., Ostmann, R., Schubert, T., Brandel, A., Richert, L., Mely, Y., Madl, J., and Römer, W. (2023). The bacterial lectin LecA from P. aeruginosa alters membrane organization by dispersing ordered domains. Commun Phys 6. 10.1038/s42005-023-01272-3.

95. Darkow, E., Rog-Zielinska, E.A., Madl, J., Brandel, A., Siukstaite, L., Omidvar, R., Kohl, P., Ravens, U., Römer, W., and Peyronnet, R. (2020). The Lectin LecA Sensitizes the Human Stretch-Activated Channel TREK-1 but Not Piezo1 and Binds Selectively to Cardiac Non-myocytes. Front. Physiol. 11. 10.3389/fphys.2020.00457.

96. Immanuel, C.N., Teng, B., Dong, B.E., Gordon, E.M., Luellen, C., Lopez, B., Harding, J., Cormier, S.A., Fitzpatrick, E.A., Schwingshackl, A., et al. (2024). Two-pore potassium channel TREK-1 (K2P2.1) regulates NLRP3 inflammasome activity in macrophages. American Journal of Physiology-Lung Cellular and Molecular Physiology 326, L367–L376. 10.1152/ajplung.00313.2023.

97. Billingham, L.K., Stoolman, J.S., Vasan, K., Rodriguez, A.E., Poor, T.A., Szibor, M., Jacobs, H.T., Reczek, C.R., Rashidi, A., Zhang, P., et al. (2022). Mitochondrial electron transport chain is necessary for NLRP3 inflammasome activation. Nat Immunol 23, 692–704. 10.1038/s41590-022-01185-3.

98. Macher, B.A., and Galili, U. (2008). The Galα1,3Galβ1,4GlcNAc-R (α-Gal) epitope: A carbohydrate of unieue evolution and clinical relevance. Biochimica et Biophysica Acta (BBA) - General Subjects 1780, 75–88. 10.1016/j.bbagen.2007.11.003.

99. Helm, J., Mereiter, S., Oliveira, T., Gattinger, A., Markovitz, D.M., Penninger, J.M., Altmann, F., and Stadlmann, J. (2024). Non-targeted N-glycome profiling reveals multiple layers of organ-specific diversity in mice. Nat Commun 15, 9725. 10.1038/s41467-024-54134-z.

100. Mestas, J., and Hughes, C.C.W. (2004). Of Mice and Not Men: Differences between Mouse and Human Immunology. The Journal of Immunology 172, 2731–2738. 10.4049/jimmunol.172.5.2731.

101. Seok, J., Warren, H.S., Cuenca, A.G., Mindrinos, M.N., Baker, H.V., Xu, W., Richards, D.R., McDonald-Smith, G.P., Gao, H., Hennessy, L., et al. (2013). Genomic responses in mouse models poorly mimic human inflammatory diseases. Proc. Natl. Acad. Sci. U.S.A. 110, 3507–3512. 10.1073/pnas.1222878110.

102. Martinon, F., Pétrilli, V., Mayor, A., Tardivel, A., and Tschopp, J. (2006). Gout-associated uric acid crystals activate the NALP3 inflammasome. Nature 440, 237–241. 10.1038/nature04516.

103. Groß, O. (2012). Measuring the Inflammasome. In Leucocytes Methods in Molecular Biology., R. B. Ashman, ed. (Humana Press), pp. 199–222. 10.1007/978-1-61779-527-5_15.

104. Mitchell, E., Houles, C., Sudakevitz, D., Wimmerova, M., Gautier, C., Pérez, S., Wu, A.M., Gilboa-Garber, N., and Imberty, A. (2002). Structural basis for oligosaccharide-mediated adhesion of Pseudomonas aeruginosa in the lungs of cystic fibrosis patients. Nat Struct Biol 9, 918–921. 10.1038/nsb865.

105. Chemani, C., Imberty, A., De Bentzmann, S., Pierre, M., Wimmerová, M., Guery, B.P., and Faure, K. (2009). Role of LecA and LecB Lectins in *Pseudomonas aeruginosa*-Induced Lung Injury and Effect of Carbohydrate Ligands. Infect Immun 77, 2065–2075. 10.1128/IAI.01204-08.

106. Schneider, K.S., Thomas, C.J., and Groß, O. (2013). Inflammasome Activation and Inhibition in Primary Murine Bone Marrow-Derived Cells, and Assays for IL-1α, IL-1β, and Caspase-1. In The Inflammasome Methods in Molecular Biology., C. M. De Nardo and E. Latz, eds. (Humana Press), pp. 117–135. 10.1007/978-1-62703-523-1_10.

107. Lagies, S., Schlimpert, M., Neumann, S., Wäldin, A., Kammerer, B., Borner, C., and Peintner, L. (2020). Cells grown in three-dimensional spheroids mirror in vivo metabolic response of epithelial cells. Commun Biol 3, 246. 10.1038/s42003-020-0973-6.

108. Lagies, S., Pichler, R., Vladimirov, G., Gawron, J., Bäzner, F., Schreiner, A., Kadena, D., Plattner, D.A., Lienkamp, S.S., and Kammerer, B. (2021). Metabolic and Lipidomic Assessment of Kidney Cells Exposed to Nephrotoxic Vancomycin Dosages. IJMS 22, 10111. 10.3390/ijms221810111.

