## Supplementary figures for "Host glycans trap bacterial lectins to trigger NLRP3 inflammasome activation"

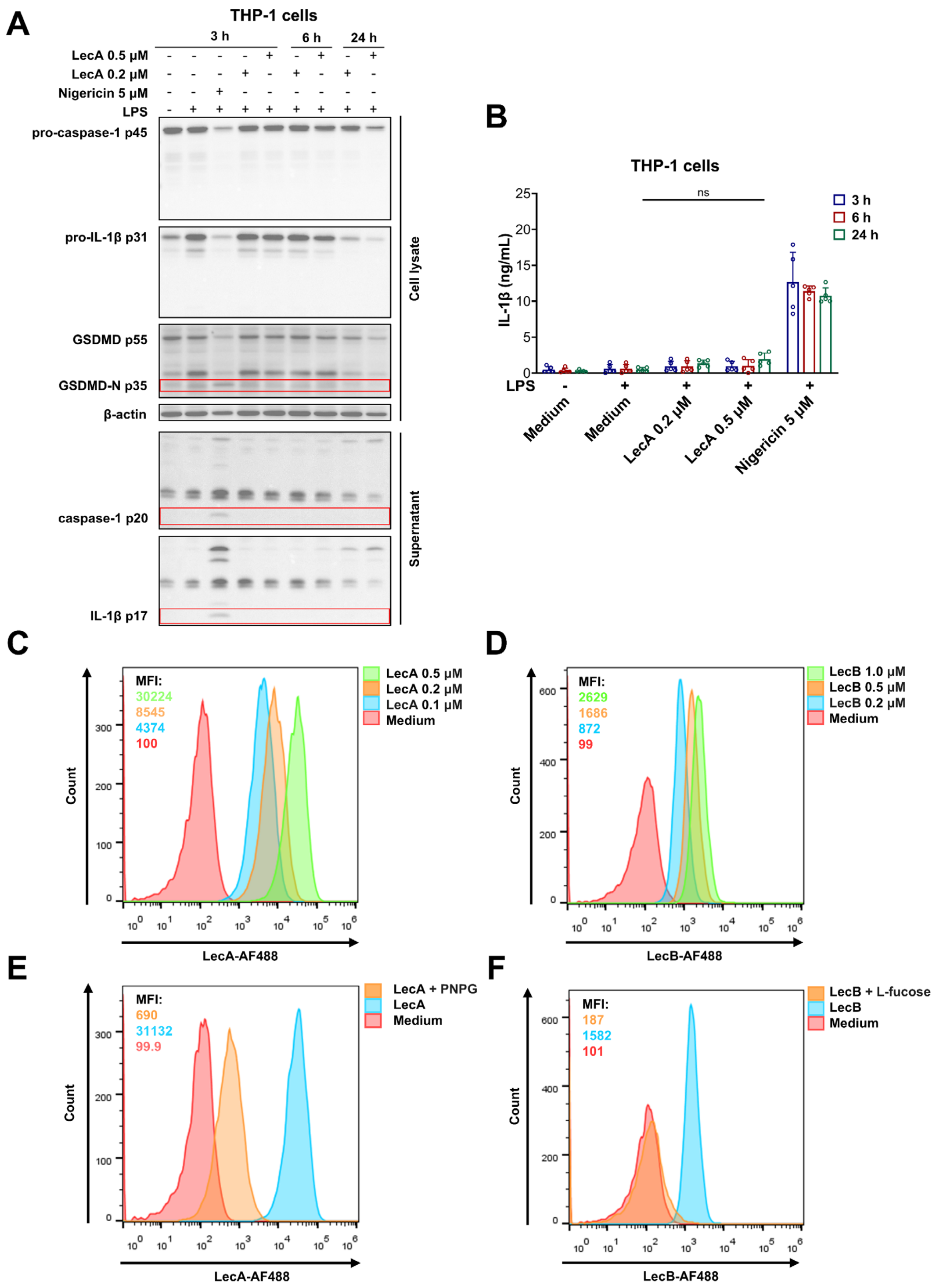


**Figure S1. LecA fails to induce inflammasome activation despite stronger binding to THP-1 cells**

(A) Immunoblot of PMA-differentiated, unprimed or LPS-primed THP-1 cells stimulated with LecA (0.2 or 0.5 µM) for 3, 6 and 24 hours or Nigericin (5 µM) for 3 hours. Red boxes indicate cleavage products. (B) ELISA for IL-1β from supernatants of THP-1 cells stimulated with LecA or Nigericin for 3, 6 and 24 hours. (C–F) Flow cytometry analysis of THP-1 cells incubated with fluorescently labeled LecA (0.1–0.5 µM; C) or LecB (0.2–1 µM; D), or with LecA or LecB

(0.5 μM each) alone or following pre-incubation with the respective sugar inhibitor PNPG (10 mM; E) or L-fucose (45 mM; F). MFI, mean fluorescence intensity. Data information: Images in (A) and flow cytometry plots in (C–F) are representative of n = 3 independent experiments. Graphs in (B) show mean ± SD from n = 5 independent experiments. MFI values in (C–F) are from the representative experiment. *P* values for indicated comparisons in (B) were determined by two-way ANOVA followed by Tukey's multiple-comparisons test. ns, not significant.

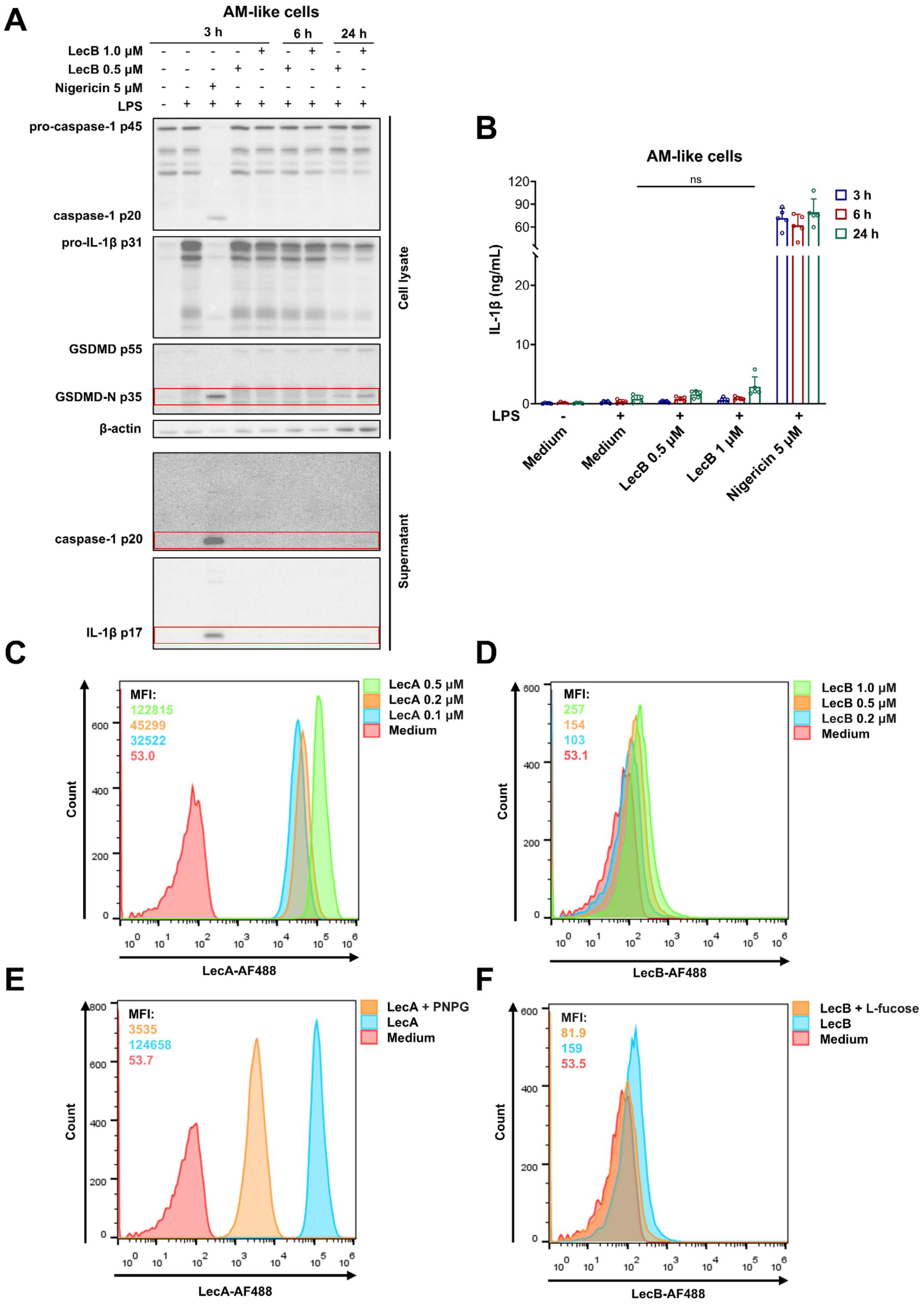


**Figure S2. LecB fails to induce inflammasome activation despite carbohydrate-dependent binding to AM-like cells**

(A) Immunoblot of unprimed or LPS-primed AM-like cells stimulated with indicated concentrations of LecB (0.2 or 0.5 µM) for 3, 6 and 24 hours or Nigericin (5 µM) for 3 hours. (B) ELISA for IL-1β from supernatants of AM-like cells stimulated with LecB or Nigericin for 3, 6 and 24 hours. (C–F) Flow cytometry analysis of AM-like cells incubated with fluorescently labeled LecA (0.1–0.5 µM; C) or LecB (0.2–1 µM; D), or with LecA or LecB (0.5 µM each) alone or following pre-incubation with the respective sugar inhibitor PNPG (10 mM; E) or L-fucose (45 mM; F). Data information: Images in (A) and flow cytometry plots in (C–F) are representative of n = 3 independent experiments. Graphs in (B) show mean ± SD from n = 5 independent experiments. MFI values in (C–F) are from the representative experiment. *P* values for indicated comparisons in (B) were determined by two-way ANOVA followed by Tukey's multiple-comparisons test. ns, not significant.

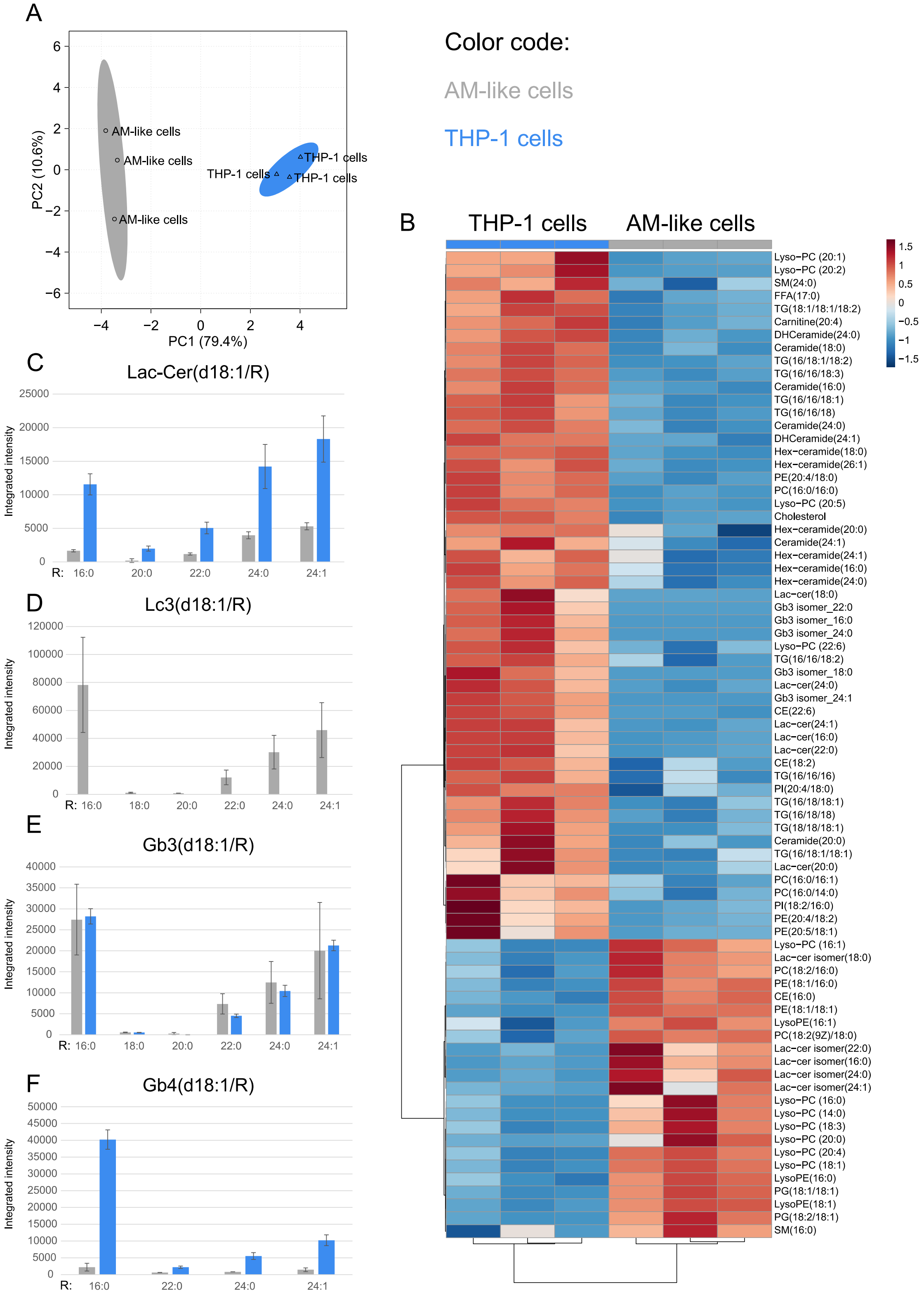


**Figure S3. Lipid profiling of murine and human macrophages**

Murine (grey) and human (blue) macrophages harbor a different lipidomic profile. A: Principal component analysis of murine AM-like cells and human THP-1 cells reveal a different lipid composition with almost 80% explained differences on PC1. B: Heatmap of significantly changed lipids after false-discovery-rate correction with $q < 0.05$. Gb3 was not significantly changed. Auto-scaled z-scores are displayed with red for higher abundance and blue for lower abundance. Lipids and samples are clustered by the Euclidian-Ward method. C–F: bar charts for $[M+H]^+$ species of lactosyl-ceramide, Lc3, Gb3 and Gb4. The "isomers" in figure B were identified as in-source fragmentation artefacts arisen from the N-acetyl-hexosylated species (Lc3 = N-acetyl-hexosylated-lactosyl-ceramide, Gb4 = N-acetyl-hexosylated-Gb3). Lc3 in panel D was not detected in THP-1 macrophages. For identification see figure S4. The fatty acyl chain (R) of each species is depicted as carbon number: double bond number. Graphs show mean ± SD from n = 3 biological replicates. Ceramides, hexosylceramides, and lactosylceramides were more abundant in THP-1 cells, whereas total Gb3 levels were comparable.

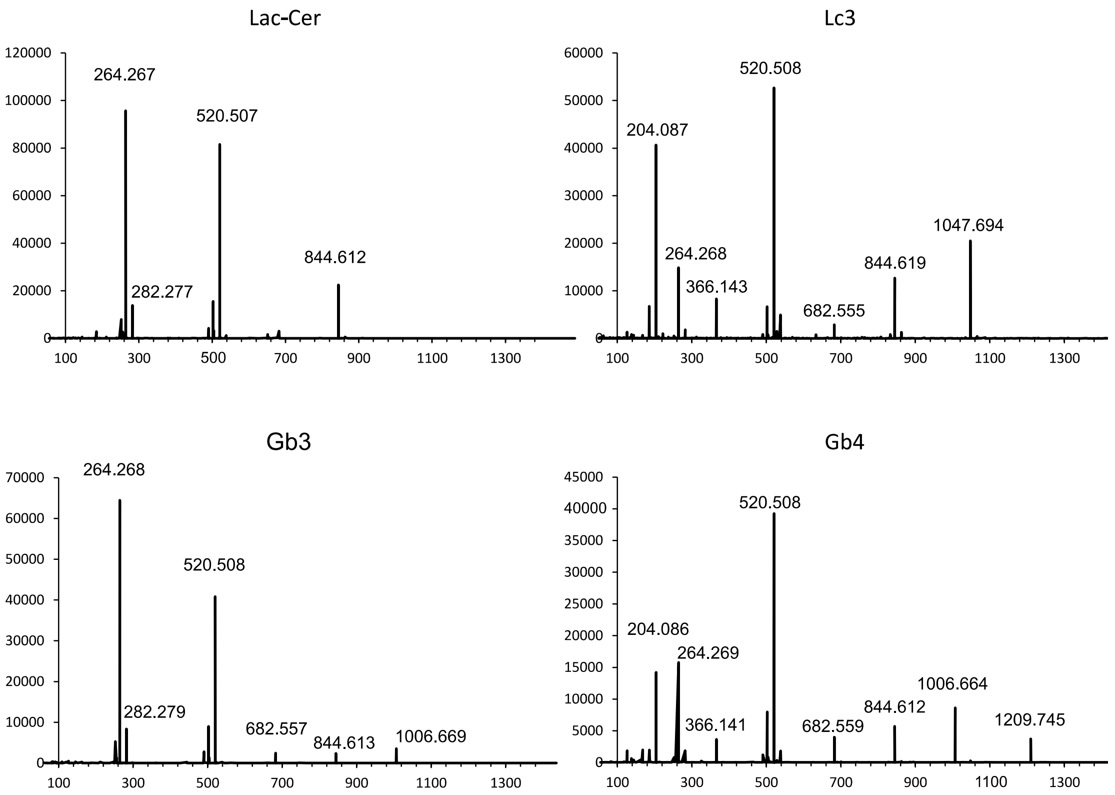


**Figure S4. Mass spectrometry of selected lipids**

Example spectra for glycosphingolipids with 16:0 acyl group for lactosyl-ceramide, Lc3, Gb3 and Gb4 in positive mode by LC-QqTOF. Spectra to structure allocations are shown in Supplementary Table S1. Multiple isomers of lactosylceramide and Gb3 were identified during peak annotation and were significantly altered between the two cell types (Figure S3B). To further resolve these species, the same samples were analyzed by untargeted LC-QqTOF. The isomers were assigned as Lc3 and Gb4. Quantification (Figure S3C–S7F) showed higher levels of lactosylceramides and Gb4 in human cells, Lc3 exclusively in murine cells, and no difference in Gb3 abundance.

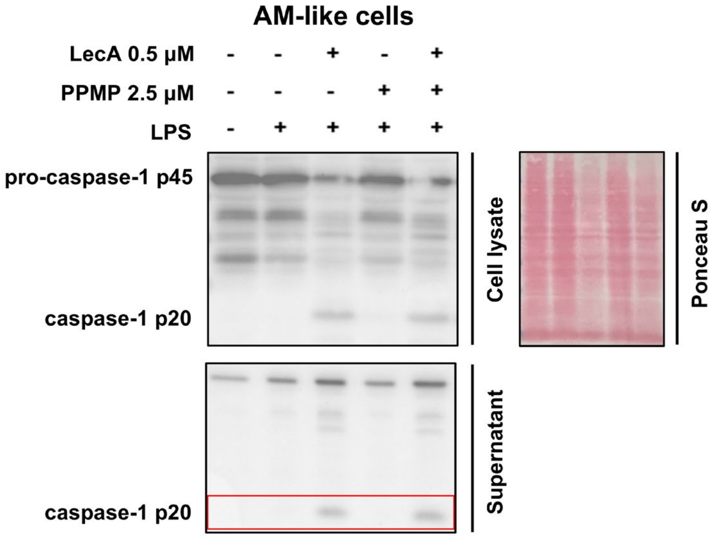


**Figure S5. Glycosphingolipid depletion has no impact on LecA-induced caspase-1 activation in AM-like cells**

Immunoblot of unprimed or LPS-primed AM-like cells stimulated with LecA (0.5 µM) for 24 hours, with or without PPMP pretreatment (2.5 µM, 4 days). PPMP was maintained throughout LecA stimulation. Ponceau S staining of the corresponding cell lysate membrane is shown.

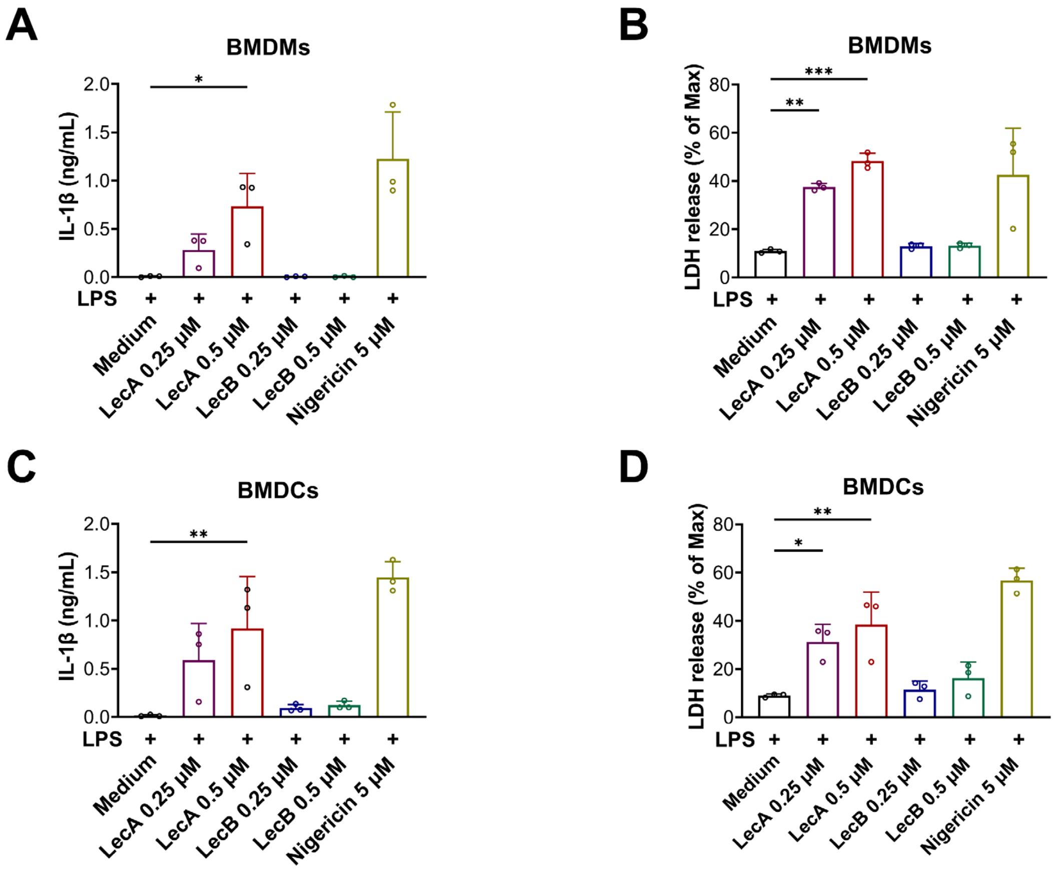


**Figure S6. LecA, but not LecB, induces IL-1β secretion and LDH release in primary murine BMDMs and BMDCs**

(A, B) IL-1β secretion (A) and LDH release (B) from primary murine BMDMs primed with LPS and subsequently stimulated with LecA or LecB (0.25 or 0.5 μM) for 24 hours or Nigericin (5 μM) for 2 hours. (C, D) IL-1β secretion (C) and LDH release (D) from primary murine BMDCs treated as in (A, B). Data information: Graphs in (A–D) show mean ± SD from n = 3 independent experiments. *P* values for indicated comparisons were determined by one-way ANOVA followed by Dunnett's multiple-comparisons test versus the LPS-only control. *$p < 0.05$; **$p < 0.01$; ***$p < 0.001$.

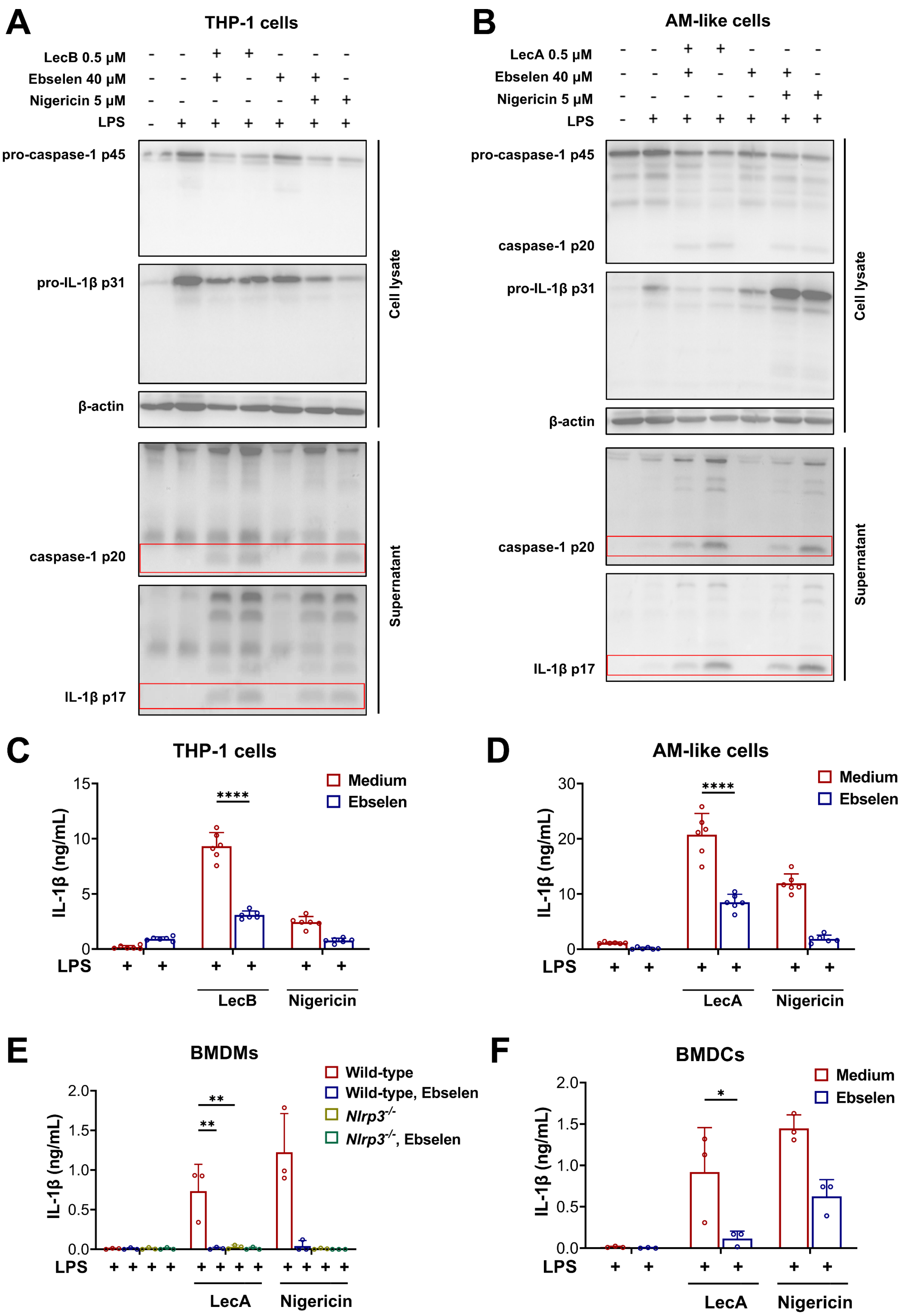


**Figure S7. Ebselen inhibits lectin-induced inflammasome activation in a cell type-dependent manner**

(A, B) Immunoblot of PMA-differentiated THP-1 cells (A) and AM-like cells (B), unprimed or LPS-primed, stimulated with LecB (0.5 µM; A) or LecA (0.5 µM; B) for 24 hours or Nigericin (5 µM) for 1 hour, with or without Ebselen (40 µM). (C, D) ELISA for IL-1β from supernatants of THP-1 cells (C) and AM-like cells (D) treated as in (A) and (B), respectively. (E, F) ELISA for IL-1β from supernatants of LPS-primed BMDMs (wild-type and *Nlrp3*$^{-/-}$; E) and BMDCs (F) stimulated with LecA (0.5 μM) for 24 hours or Nigericin (5 μM) for 2 hours, with or without Ebselen (30 μM). Data information: Images in (A, B) are representative of n = 3 independent experiments. Graphs in (C, D) show mean ± SD from n = 6 independent experiments. Graphs in (E, F) show mean ± SD from n = 3 independent experiments. *P* values for indicated comparisons in (C–F) were determined by two-way ANOVA followed by Tukey's multiple-comparisons test. *$p < 0.05$; **$p < 0.01$; ****$p < 0.0001$.

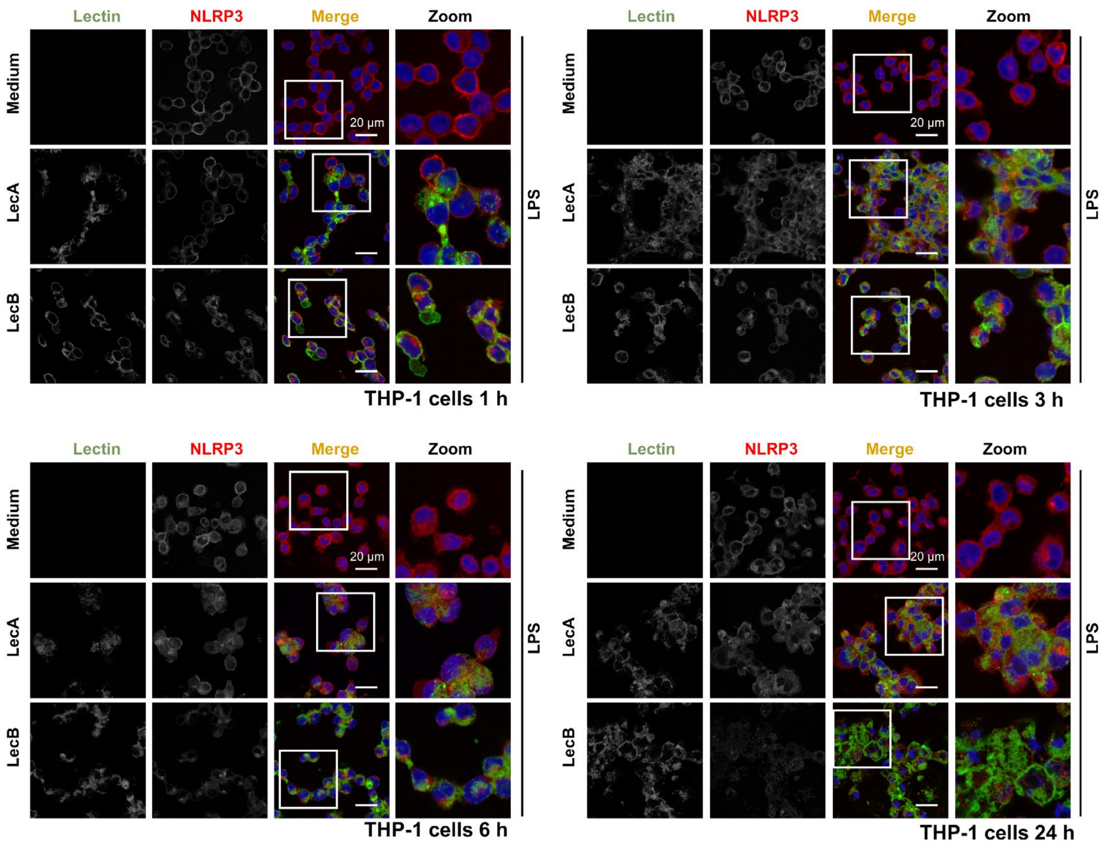


**Figure S8. LecB, but not LecA, stimulates NLRP3 relocalization in THP-1 cells**

Representative confocal images of PMA-differentiated, LPS-primed THP-1 cells incubated with LecA-AF488 or LecB-AF488 (0.5 µM) for 1, 3, 6 and 24 hours to assess lectin binding and NLRP3 relocalization. Cells were stained for NLRP3 (red) and nuclei (DAPI, blue), and lectin binding is shown in green. Representative images from multiple fields of view are shown. Scale bars, 20 µm.

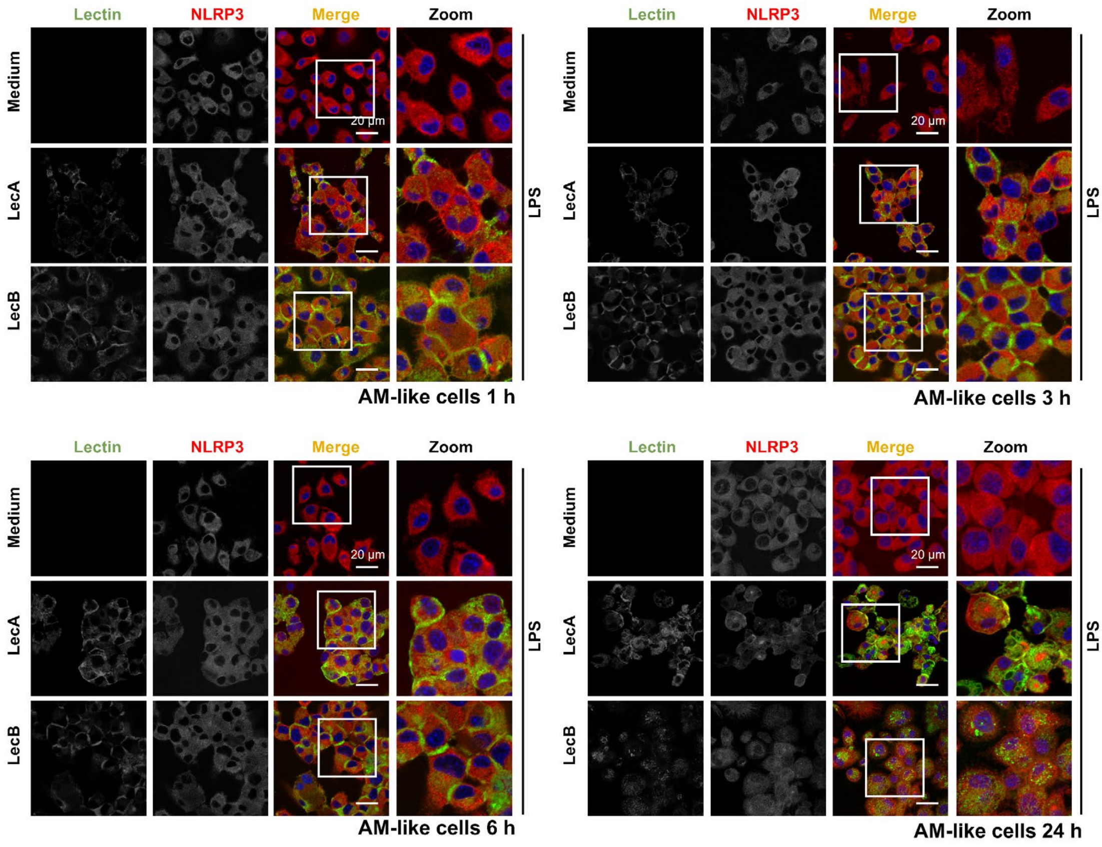


**Figure S9. LecA, but not LecB, stimulates NLRP3 relocalization in AM-like cells**
Representative confocal images of LPS-primed AM-like cells incubated with LecA-AF488 or LecB-AF488 (0.5 µM) for 1, 3, 6 and 24 hours to assess lectin binding and NLRP3 relocalization. Cells were stained for NLRP3 (red) and nuclei (DAPI, blue), and lectin binding is shown in green. Representative images from multiple fields of view are shown. Scale bars, 20 µm.

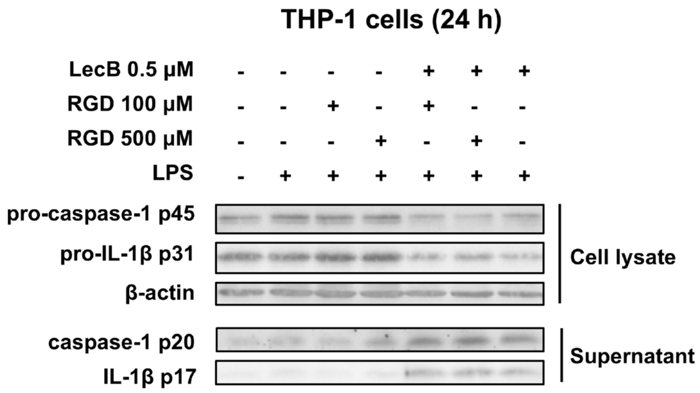


**Figure S10. RGD peptide fails to inhibit LecB-induced inflammasome activation in human THP-1 cells**

Immunoblot of PMA-differentiated, unprimed or LPS-primed THP-1 cells stimulated with LecB (0.5 µM) for 24 hours, with or without RGD peptide pretreatment (100 or 500 µM). Data information: Images are representative of n = 3 independent experiments.

**Supplementary Table S1.**

Fragment allocation. Example spectra with explanation of fragments obtained by LC-QqTOF $MS^2$ analysis. $[M+H]^+$: precursor, with sequential neutral losses (Hex: hexoside, N-Ac-Hex: N-acetylated hexosamine, R (16:0): hexadecyl) with the measured *m*/*z* values. The underlined fragments are the same fragments as in the multiple reaction monitoring analysis by LC-QqQ, explaining why the in-source fragment of Lc3 and Gb4 were identified as Lac-Cer and Gb3 isomers. Lac-Cer and Gb3 retention times were validated by authentic standards. Lc3 and Gb4 are only identified by MS/MS experiments and they could also be an isomer like ganglioside GA2 instead of Lc3 or isoGb4 instead of Gb4. Lc3 and Gb4 were chosen based on their described role in macrophage biology. [1]

| **Fragment** | **Lac-Cer (d18:1/16:0)** | **Lc3 (d18:1/16:0)** | **Gb3 (d18:1/16:0)** | **Gb4 (d18:1/16:0)** |
|---|---|---|---|---|
| | *m*/*z* | *m*/*z* | *m*/*z* | *m*/*z* |
| **$[M+H]^+$** | 862.625 | 1065.709 | 1024.683 | 1227.764 |
| **-H2O** | 844.612 | 1047.694 | 1006.669 | 1209.745 |
| **-N-Ac-Hex** | NA | 844.619 | NA | 1006.664 |
| **-Hex** | 682.560 | 682.555 | 844.613 | 844.612 |
| **-Hex** | 520.507 | 520.508 | 682.557 | 682.559 |
| **-Hex** | NA | NA | 520.508 | 520.508 |
| **-R (16:0)** | 264.267 | 264.268 | 264.268 | 264.269 |
| **$[N\text{-}Ac\text{-}Hex2+H]^+$** | NA | 366.146 | NA | 366.141 |
| **$[N\text{-}Ac\text{-}Hex+H]^+$** | NA | 204.087 | NA | 204.086 |

[1] https://www.frontiersin.org/journals/immunology/articles/10.3389/fimmu.2019.00090/full
